# Effects of environmental change on population growth: monitoring time-varying carrying capacity in free-ranging spotted hyenas

**DOI:** 10.1101/2024.04.11.589105

**Authors:** Liam D. Bailey, Oliver P. Höner, Eve Davidian, Arjun Dheer, Viktoriia Radchuk, Leonie F. Walter, Ella W. White, Alexandre Courtiol

## Abstract

In conservation, a growing population is often taken as a sign of success. But trends in population size can be misleading. When individuals are long-lived, populations may keep growing—for a time—even as the environment begins to stabilize or deteriorate. Trends in carrying capacity (*K*) would better reflect the situation that a population finds itself in, yet *K* is commonly assumed to be static. We developed an individual-based modelling approach to estimate time-varying carrying capacity (*K*_t_) in a population of free-ranging spotted hyenas. *K*_t_ noticeably varied due to events such as disease outbreaks, but not in response to recent conservation interventions. Although the population tripled in size during the 26-year study period, we found no corresponding trend in *K*_t_. Rather, population recovery resulted from earlier environmental improvements. Our approach, which can be applied to populations of other species, allows for a faster and more accurate assessment of conservation needs.

## 1 **Introduction**

The world is confronting an acute biodiversity crisis, with population declines and extinctions occurring across major taxonomic groups (Barnosky et al., 2011; Ceballos et al., 2017; Pimm et al., 2014; WWF, 2022). Changes in the biotic and abiotic environment, such as the expansion of invasive species, habitat loss, and climate change, are key factors driving modern biodiversity loss (Mazor et al., 2018). Consequently, many conservation programs strive to improve environmental conditions to achieve species conservation. For example, conservation programs may protect habitats from human disturbance (Maxwell et al., 2020), reduce competition from invasive species (Prior et al., 2018), or increase the availability of limited resources such as food and shelter (Lindenmayer et al., 2009; Murray et al., 2016). In contrast, the evaluation of conservation outcomes often relies on measuring trends in population size rather than changes in environmental conditions (Badalotti et al., 2022; Crees et al., 2016; IUCN, 2012; Mace et al., 2008; WWF, 2022). Unfortunately, population size is an unreliable indicator of recent changes in environmental conditions, potentially undermining the suitability of conservation decisions.

Both population growth and decline can occur independently of recent trends in environmental conditions (Fig. 1). In some cases, such as in short-lived and highly fecund species, population growth can occur rapidly in response to improving environmental conditions (Fig. 1A; e.g., Monaghan et al., 2020). However, the maximum attainable population growth rate is restricted by the structure of the population (e.g., sex ratio and age distribution) and species-specific life-history constraints (e.g., maximum litter size; Ackleh et al., 2018; Coulson and Hudson, 2002; Koons et al., 2007). As a consequence, populations cannot always grow as rapidly as the environment changes. Trends in population size reflect the combined effect of environmental change that has occurred in the past and not recent environmental change alone.

**Figure 1:**
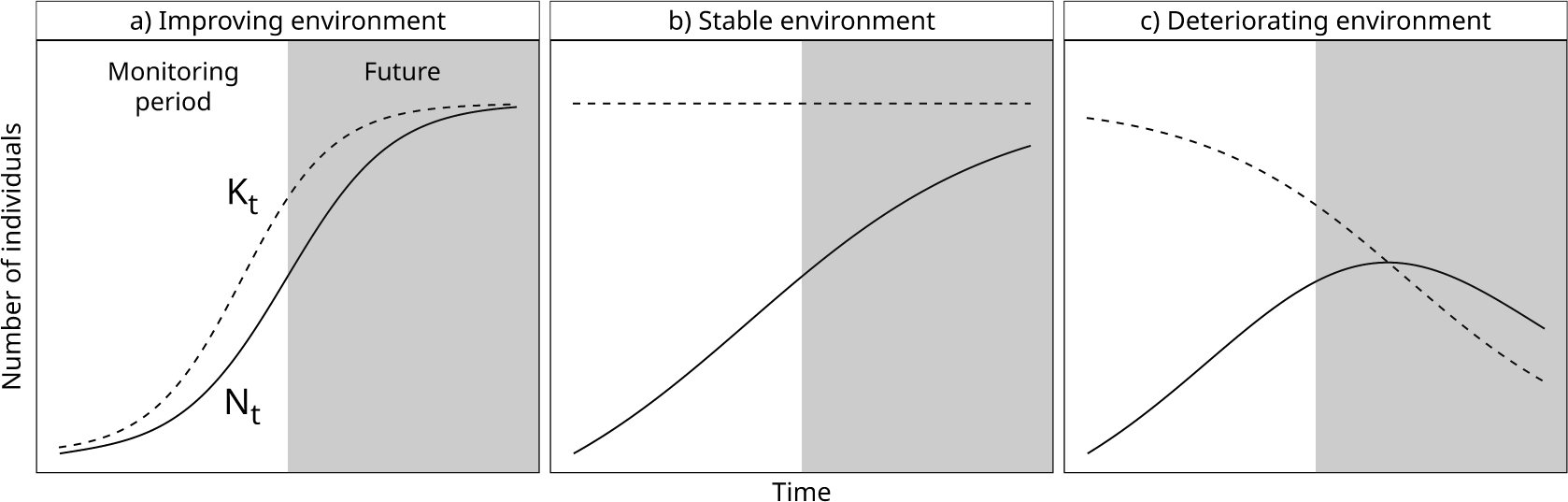
Population growth can occur under drastically different scenarios of environmental change. Plots show changes in population size (*N*_t_; solid line) and time-varying carrying capacity (*K*_t_; dashed line) under three hypothetical scenarios. Population monitoring is ongoing in the white region (left). The grey shaded region (right) shows future changes. Population size at the beginning or end of the monitoring period is identical across all three panels. a) Improving environment: The population is near carrying capacity at the start of the monitoring period. Carrying capacity increases (e.g., due to successful conservation) and population size increases rapidly in response to change in *K*_t_. b) Stable environment: Carrying capacity increases before the monitoring period but remains constant thereafter. Population growth rate responds slowly to changes in carrying capacity and the population remains well below carrying capacity during much of the monitoring period (i.e., *N*_t_ *≪ K*_t_). c) Deteriorating environment: Similar to scenario b), population size starts well below carrying capacity (i.e., *N*_t_ *≪ K*_t_); however, during the monitoring period carrying capacity declines. Deterioration in environmental conditions eventually leads to population decline once *N*_t_ *> K*_t_, but this can happen after the monitoring period has ended. The three scenarios were generated assuming a simple discrete time logistic growth model. In scenario a) and c), carrying capacity is also itself a logistic function of time (see Meyer and Ausubel, 1999).

Limits on population growth are particularly important in species with longer generation times, such as long-lived mammals (Coulson and Hudson, 2002). In such circumstances, population growth may persist even after environmental conditions have stabilised (Fig. 1B). In fact, lingering effects of old environmental improvement can mask recent declines in environmental conditions (Fig. 1C). To complicate matters further, changes in population size can also occur due to non-environmental processes such as demographic stochasticity (Lande, 1993) or positive density dependence (i.e., Allee effects; Courchamp et al., 1999; Stephens et al., 1999). Therefore, trends in population size cannot be trusted to detect recent trends in environmental conditions. Alternative metrics directly derived from population size (e.g., instantaneous growth rate; Sibly and Hone, 2002) have similar limitations.

To avoid the problems caused by a reliance on population size, conservation assessments should also attempt to measure changes in the biotic and abiotic environment over time. *Carrying capacity* (often referred to as *K*) is one metric that can be adapted for this purpose. This metric is indeed commonly used in conservation to capture the effects of all environmental variables that influence a target population, even those that may be hard, or impossible, to measure directly. *K* is often treated as a fixed number of individuals that could be supported by the environment (Chapman and Byron, 2018; Dhondt, 1988). However, the environment, and therefore carrying capacity, both vary over time. Variation in the environment necessarily influences the size of a population through vital rates (here defined as functions of age and other characteristics of an individual defining its survival and fertility). Therefore, carrying capacity can be understood as an emergent property of the vital rates of all the individuals of a population. Rather than treating carrying capacity as static as it is usually done, it is therefore useful to consider carrying capacity as variable (e.g., Britten et al., 2017; Fukasawa et al., 2013; Hinrichsen and Paulsen, 2020; Kaeriyama et al., 2009; Meyer and Ausubel, 1999; Zehnder and Hunter, 2008). For clarity, we refer to carrying capacity that is allowed to vary over time as *time-varying carrying capacity* or *K*_t_.

In this paper, we introduce a method to quantify *K*_t_ using a combination of vital rate estimation and population projection performed using individual-based modelling (DeAngelis and Grimm, 2014; Judson, 1994) (see supplementary section S1.1 & S1.2 ; Table S1; Fig. S1–S5). To illustrate our method, we apply it to a free-ranging population of large carnivores: the spotted hyenas (*Crocuta crocuta*) of Ngorongoro Crater, Tanzania (see section 4.1 in Methods for details on the study species and population). This population has been monitored continually for almost three decades (April 1996–present), making it one of the most closely monitored large carnivore populations in the world (Bonnet et al., 2022). For much of the first two decades of study, spotted hyenas in Ngorongoro Crater experienced population growth (Höner et al., 2005, 2012). This population growth could be attributed to conservation interventions and natural phenomena that have occurred during the study period (Table ED1), such as increased restrictions on road use or changes in spotted hyena prey abundance (Höner et al., 2005). However, as we have discussed above, such population growth could also be the consequence of the lingering effects of environmental change that occurred before monitoring began. Quantifying *K*_t_ helps to disentangle these possible explanations.

In our analysis, we quantified time trends in population size, vital rates, and *K*_t_ in the focal population of spotted hyenas between 1997 and 2022. We used *K*_t_ to estimate environmental conditions in each year and assess whether population growth indicates a recent improvement in environmental conditions or lingering effects of older environmental change. We expected physiological constraints on reproduction in spotted hyenas (see section 4.1) to place marked limits on maximum possible population growth, making observed population growth likely to be affected by environmental change that occurred several years before present. Estimating *K*_t_ also allowed us to determine whether the population has reached demographic equilibrium, and to assess how different variables (e.g., disease, competition) have driven carrying capacity over the past 26 years. In the discussion, we detail how similar assessments could be performed in other populations or species.

## 2 **Results**

### 2.1 Population size and demographic composition

In the late 1960s, Kruuk (1972) assessed the number of adult spotted hyenas in Ngorongoro Crater at 385 individuals. Between the 1960s and 1990s, the population experienced a large decline due to undocumented reasons. When the long-term monitoring of the population began (April 1996), population size (*N*_t_) was down to 31% of the 1960s estimate (119 adults, 65 juveniles). Monitoring since April 1996 documents significant population growth across the study period (Fig. 2; *β_N_*_1997_*_−_*_2022_ = 13.4; 95% Confidence Interval or CI_95%_ = 0.587/24.8; Likelihood Ratio Test: *χ*^2^ = 4.09; df = 1; *p* = 0.043). From 1997 to 2011, population size grew at a rate of 31 individuals per year (*β_N_*_1997_*_−_*_2011_ = 30.8; CI_95%_ = 23.5/39.7; Likelihood Ratio Test: *χ*^2^ = 21.0; df = 1; *p <* 0.001). In September 2011, the numbers of adults in Ngorongoro Crater returned, for the first time since April 1996, to numbers similar to Kruuk’s estimates from the 1960s (378 adults, 185 juveniles). No significant time-trend in population size occurred after 2011 (*β_N_* = -9.97; CI_95%_ = -31.7/12.2; Likelihood Ratio Test: *χ*^2^ = 1.10; df = 1; *p* = 0.29), although the annual population size varied around a median of 589 individuals (range: 454– 712), with periods of sharp decline recorded in 2012–2013, 2018–2019 and 2019–2020. The most recent population decline occurred during an outbreak of *Streptococcus* infection, with the population reaching in May 2020 its lowest size since 2012 (354 individuals). Despite large changes in population size, the age and sex composition of the population remained relatively stable during the entire study period (see section S2.1; Fig. S6).

**Figure 2:**
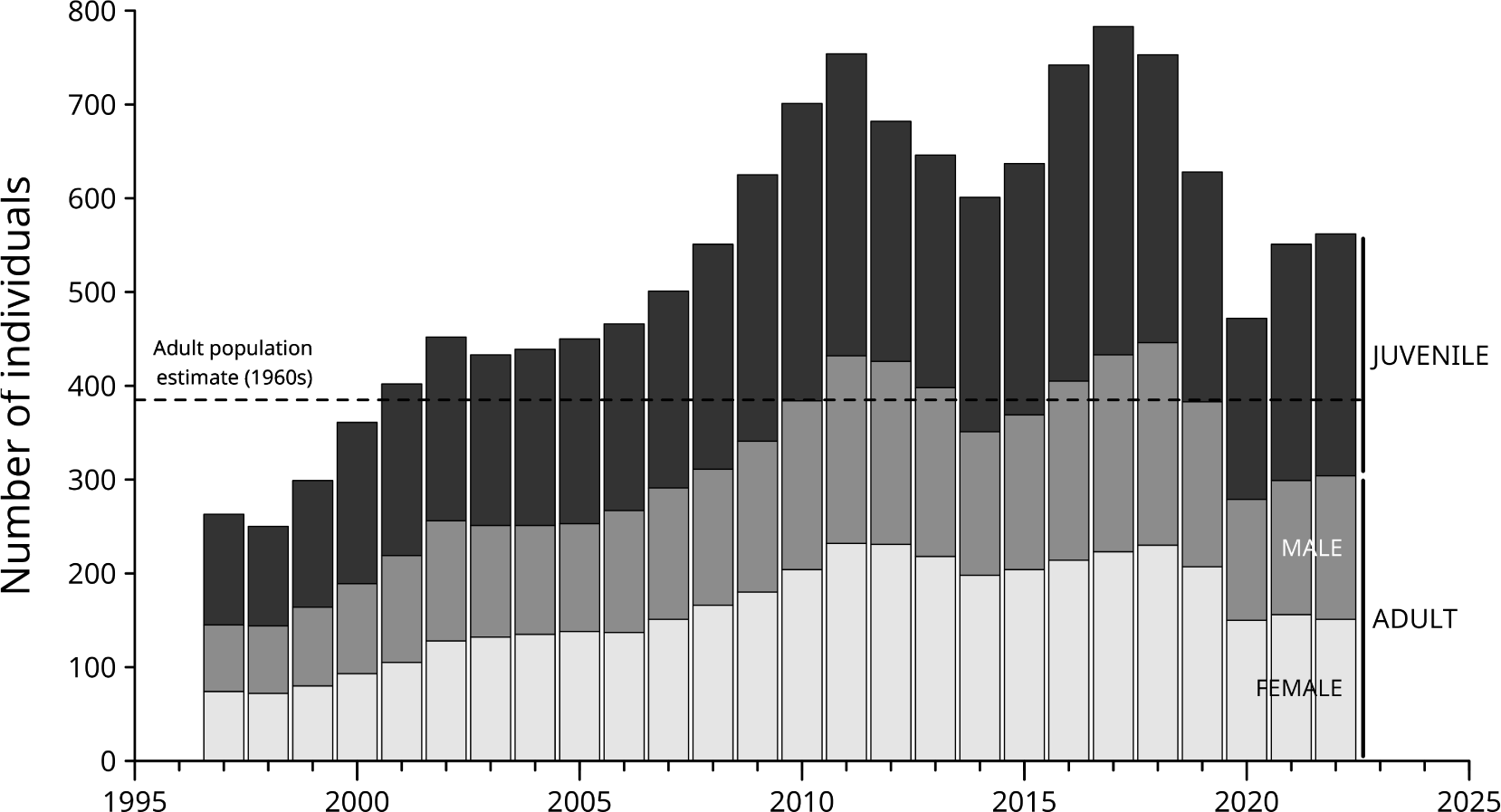
Annual population size of spotted hyenas in Ngorongoro Crater, Tanzania since 1996. Colours distinguish adults (*≥* 24 months) of known sex (female: light grey; male: dark grey) and juveniles (darkest grey). The stacked areas indicate the contribution of each subgroup to the entire population size, such that the total population size with adults and juveniles can be read from the top of the stack. Seven adults with unknown sex were excluded. Horizontal dashed line represents estimated number of adult individuals during the 1960s (Kruuk, 1972).

### 2.2 Carrying capacity

To assess changes in environmental conditions, we estimated time-varying carrying capacity (*K*_t_), for each year from 1997 to 2022, using an individual-based modelling approach (see Fig. ED1; section S1.1). Carrying capacity was not estimated in 1996 and 2023 as demographic information was too uncertain in these first and last years of monitoring. Carrying capacity in each year (t) was estimated as the asymptotic population size around which our individual-based model stabilised when conditions observed in year t were assumed to persist. Our individual-based model relies on a set of statistical models fitted on vital rates using the long-term data collected on spotted hyenas in Ngorongoro Crater (Table ED2; section S1.1.7). These models, fitted with the R package spaMM (Rousset and Ferdy, 2014), were used to simulate the reproduction and survival of individuals at each time step of the simulation (months). Importantly, we did not directly impose an artificial upper limit on population size as is the case in the logistic model of population growth found in ecology textbooks (see Mallet, 2012). Instead, the statistical models consider that density dependence impacts vital rates (Fig. ED2; section S2.2; Fig. S7–S9). As mentioned in introduction, *K*_t_ is thus an emergent property of the individual-based model (Grimm and Railsback, 2005; Radchuk et al., 2019). We assumed that density dependence operates at the level of spotted hyena social groups (clans), meaning that the size of each of the eight clans in a simulation run stabilises around an asymptotic clan size. Our approach thus estimates a value of carrying capacity for each clan individually (denoted *K*_tc_) as well as the population as a whole 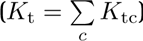 The statistical models of vital rates also include a random factor describing environmental variation in time (year) and space (clan). The difference in random effect realizations associated with each year-clan combination is the cause of all variation in carrying capacity between years in our simulations.

In contrast to population size which increased over years during much of the monitoring period, we identified no linear trend in *K*_t_ (Fig. 3; *β_K_*_1997−2022_ = -5.24; CI_95%_ = -11.92/1.16; Likelihood Ratio Test: *χ*^2^ = 2.21; df = 1; *p* = 0.14). Across the entire study (1997–2022), the median *K*_t_ value was 562 individuals (arithmetic mean = 590; harmonic mean = 552; see section S2.3 for details; Fig. S10). Our computations revealed three periods of sharp environmental decline where *K*_t_ decreased to *≤* 80% of the previous year (2001–2002, 2010–2011, 2017–2019). None of these periods match any of the periods of sharp population size decline mentioned above. At the clan level, one of the eight clans did experience a significant linear change in *K*_tc_ with time (Table S2), but the trend was decreasing and not increasing (Lemala clan: *β*_year_ = -2.57; CI_95%_ = -4.50/-0.526; section S2.4). We also identified no significant trend in either *K*_t_ or *K*_tc_ when focusing on the period of population growth (1997–2011; section S2.4; Table S3).

**Figure 3:**
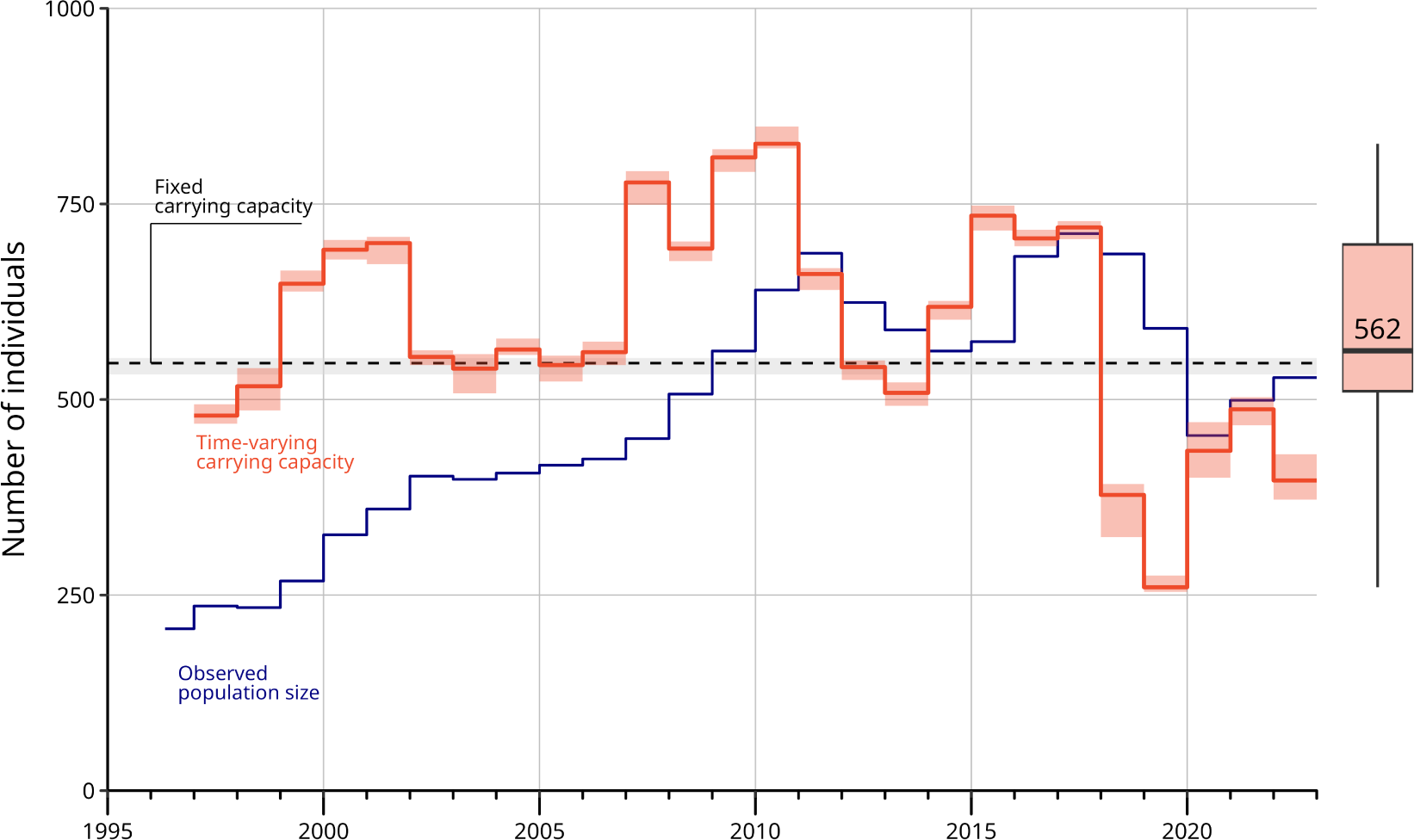
Carrying capacity of spotted hyenas shows large inter-annual variation but no linear trend over the years. The solid red line shows annual median carrying capacity estimated using the ‘Spotted Hyena Individual-based Model’ (SHIM). The marginal boxplot shows the distribution of median carrying capacity values across all years (1997–2022). The median value of all annual estimates was 562. The dashed line shows fixed carrying capacity estimated from SHIM without considering inter-annual variation, which closely matches estimates using traditional demographic models (section S2.3). The shaded areas around carrying capacity values show the range of carrying capacity estimates across ten simulation runs. The blue solid line shows annual population size of all spotted hyenas in Ngorongoro Crater (juveniles and adults of both sexes) since April 1996. Estimates of population size and time-carrying capacity are less reliable for the last year analysed (2022) and should thus be interpreted with caution. This is because occurrences of births and deaths rely, in part, on parentage analyses and re-sightings that are still ongoing at the time of writing.

### 2.3 Population size and carrying capacity

The comparison of inter-annual trends in population size and carrying capacity reveals that the recorded period of population growth (1997–2011) is likely to qualify as a genuine population recovery. Indeed, not only *N*_t_ recovered to its 1960s level in 2011, but this happened while *K*_t_ remains relatively stable. When the study period began, *N*_t_ was more than 200 individuals below carrying capacity when *K*_t_. *N*_t_ grew almost continuously between 1997 and 2011 but remained more than 100 individuals below *K*_t_ until 2010.

The results also show that trends in population size often failed to reflect recent change in the underlying environment. Inter-annual change in carrying capacity (*λ_K_*= *K*_t+1_*/K*_t_) and population size (*λ_N_* = *N*_t+1_*/N*_t_) were not significantly correlated (Pearson’s correlation = -0.0979; *p* = 0.64; Spearman correlation = 0.102; *p* = 0.63). We also observed all four possible combinations between the increase or decrease in either population size or carrying capacity (Fig. 4). For example, from 2006 to 2007, both carrying capacity and population size increased. In contrast, between 2001 and 2002, population size increased even though carrying capacity was decreasing. From 2018 to 2019, both population size and carrying capacity declined concurrently. Finally, between 2019 and 2020, population size declined although carrying capacity increased. Examples of population decline, such as 2018–2019 and 2019–2020, occurred when population size was above estimated *K*_t_ (Fig. ED3; Fig. S11). We never observed *N*_t_ to decline when *N*_t_ *≪ K*_t_, nor *N*_t_ to grow when *N*_t_ *≫ K*_t_ (section S2.5; Fig. ED3; Fig. S11), which supports the validity of our demographic model.

**Figure 4:**
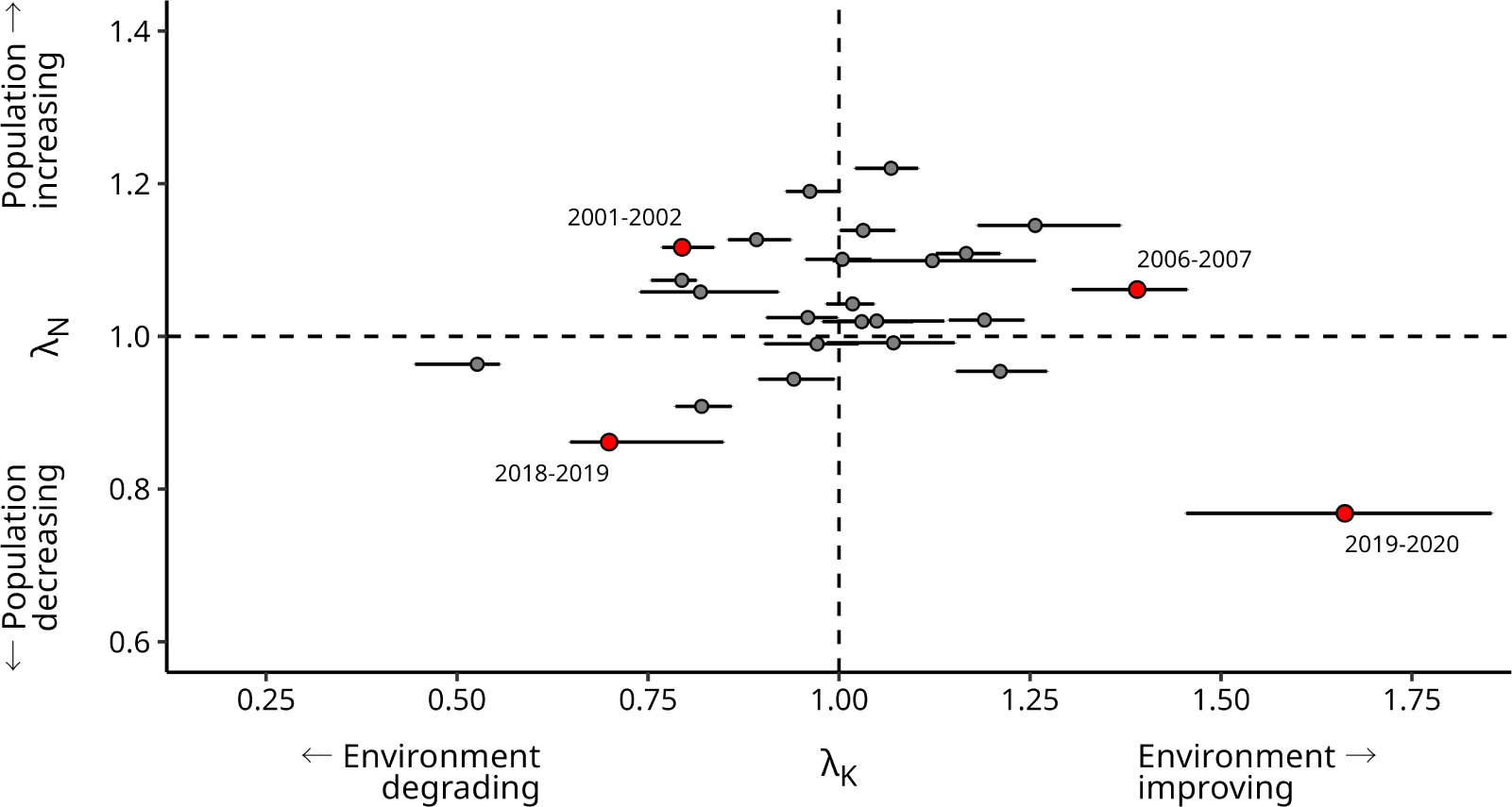
Inter-annual change in population size (*λ_N_*) is uncorrelated with inter-annual change in time-varying carrying capacity (*λ_K_*) in spotted hyenas. The points represent median values of *λ*. Red points highlight the four different examples discussed in the text (2001–2002, 2006–2007, 2018–2019, 2019–2020). The horizontal error bars represent uncertainty in *λ_K_*and show the range of possible *λ_K_* values calculated from all possible combinations of simulation runs between two focal years (*K*_t_ was estimated 10 times for each year). See Fig. S11 for plot with year labels for all points.

### 2.4 Environmental drivers and carrying capacity

While we observed no significant linear trend in *K*_t_ between 1997 and 2022 (Fig. 3), nor significant trend in vital rates that would match with the recorded trends in *N*_t_ (Fig. ED4; section S2.6; Table S4; Fig. S12), fluctuations in vital rates over time (and especially in female survival; section S2.7) did translate into large inter-annual variation in *K*_t_ (range = 260–827; *σ* = 140). The presence of such inter-annual variation in *K*_t_ provided us with the opportunity to explore the effect of external drivers (i.e., abiotic and biotic environmental variables) that may impact carrying capacity. We fitted a statistical model predicting carrying capacity at the clan level (*K*_tc_) based on five potential candidate predictors. Predictors were selected *a priori* so as to capture major potential drivers of population change (inter-specific interactions, infectious disease and anthropogenic impacts/conservation measures). We conducted analysis at the clan level to account for variation in predictors expected between clans (e.g., heterogeneity in abundance of prey and competitors). We included a semi-quantitative estimation of prey abundance and a quantitative estimation of the abundance of lions (*Panthera leo*)—the main competitor of spotted hyenas—as two variables capturing the impact of inter-specific interactions. We included the proportion of spotted hyenas observed with signs of infectious disease as a quantitative variable to account for effects of pathogens. We also included two binary variables characterising known conservation measures—whether road use was restricted and whether Maasai pastoralists were allowed to visit Ngorongoro Crater with cattle (Table ED1). All quantitative variables were Z-scored for facilitating the comparison between estimates.

Carrying capacity was significantly impacted by prey abundance, infectious disease, and lion competitors (Fig. 5; section S2.8; Table S5). As could be expected, clan territories were able to support more individuals when there was higher prey abundance (*β*_prey_ = 6.71; CI_95%_ = 3.59/10.6; Likelihood Ratio Test: *χ*^2^ = 11.8; df = 1; *p <* 0.001). Furthermore, carrying capacity was significantly negatively effected by infectious disease (*β*_disease_ = -4.47; CI_95%_ = - 7.77/-0.818; Likelihood Ratio Test: *χ*^2^ = 6.00; df = 1; *p* = 0.014) and lion density (*β*_lion_ = -6.30; CI_95%_ = -10.5/-1.22; Likelihood Ratio Test: *χ*^2^ = 6.43; df = 1; *p* = 0.011). In contrast, management interventions to restrict road use (*β*_road use inactive_ = -17.1; CI_95%_ = -32.9/3.94; Likelihood Ratio Test: *χ*^2^ = 1.07; df = 1; *p* = 0.30) and to regulate cattle grazing in Ngorongoro Crater (*β*_grazing present_ = -16.2; CI_95%_ = -31.8/2.53; Likelihood Ratio Test: *χ*^2^ = 3.23; df = 1; *p* = 0.072) had no significant effect on carrying capacity estimates.

**Figure 5:**
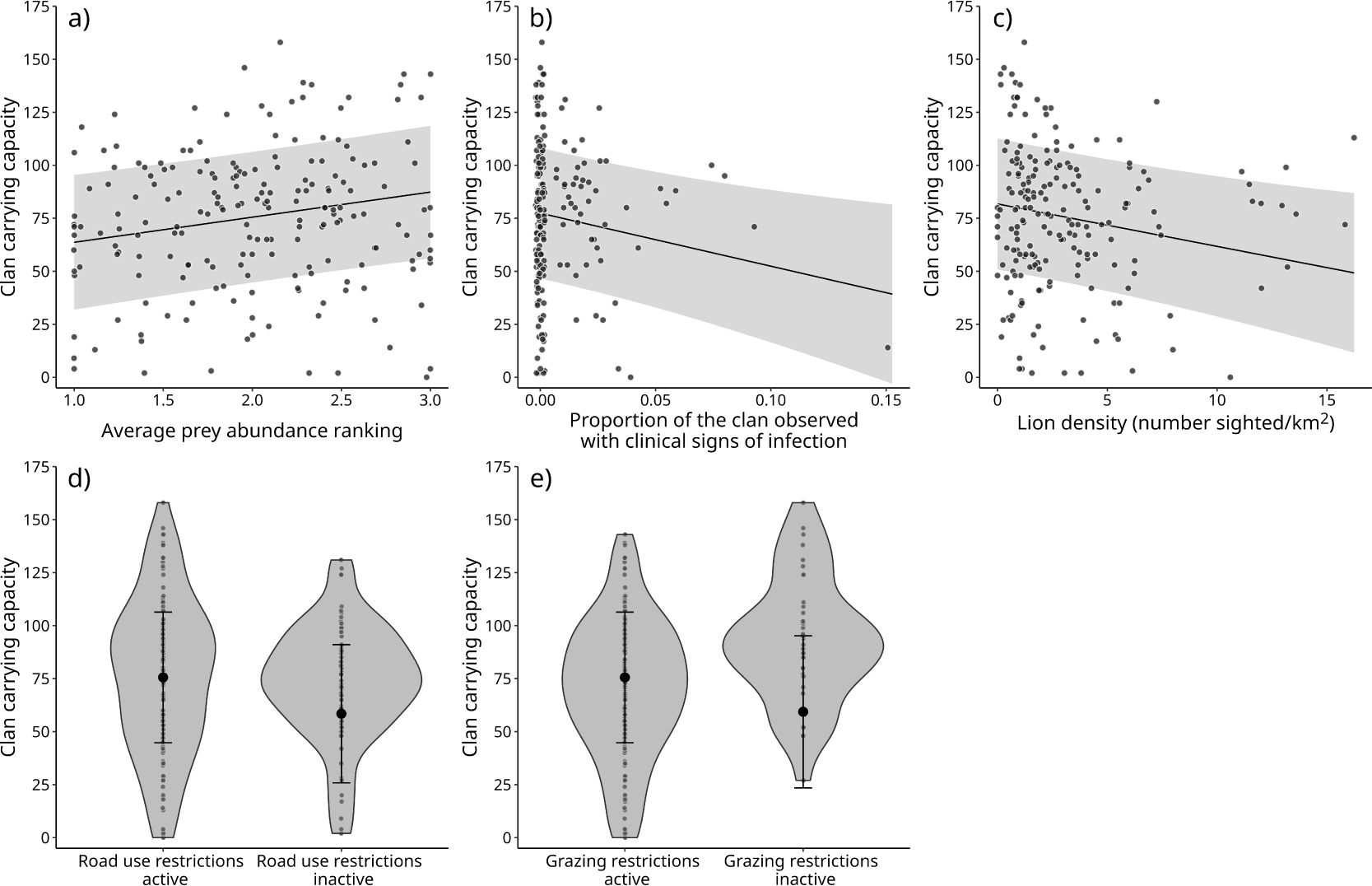
Carrying capacity was negatively affected by disease prevalence and positively affected by prey. Plotted predictions show changes in carrying capacity given changes in a) prey abundance, b) proportion of infectious disease sightings, c) lion density, d) road use restrictions, and e) grazing restrictions. Note that points on plots a-c are slightly jittered by 0.002 units on the x-axis to make them easier to distinguish. In plots d-e, grey shaded violin and transparent points represent observed data, while black solid point and error bar represent plotted predictions. All predictions were computed, for each value sampled along the x-axis as the average of predicted values on the response scale, over the empirical distribution of all other fixed-effect variables and of inferred random effects (i.e., as partial dependence effects as defined in the R package spaMM). This method allows for the visualisation of the effect of specific predictor variables while avoiding any effects caused by association with other predictor variables.

## 3 Discussion

Modern conservation places a strong focus on monitoring trends in population size (IUCN, 2012; WWF, 2022), but this approach has its limitations because population size is an unreliable indicator of recent environmental conditions. In this paper, we show how time-varying carrying capacity (*K*_t_) can be used to more directly estimate environmental conditions and provide additional information to that gained from population size. We apply our approach to study a free-ranging population of spotted hyenas in Ngorongoro Crater, Tanzania using an individual-based model—the ‘Spotted Hyena Individual-based Model’ (SHIM)—parameterised on 26 years of long-term data (1997–2022). *K*_t_ showed large inter-annual variability, likely as a consequence of changes in prey abundance, infectious disease outbreaks, and competition with lions. However, in contrast to the trajectory of population size which showed a near continuous recovery during the first 15 years of the study period (i.e., 1997–2011), carrying capacity showed no significant linear trend over time. We are now going to discuss what these findings can tell us about the current demographic status of the population and the efficacy of past and future conservation interventions. We will then discuss how our approach could be applied to other wildlife populations.

When monitoring began in 1996, the population size of spotted hyenas in Ngorongoro Crater was substantially smaller than in the late 1960s (Höner et al., 2005; Kruuk, 1972). Our estimates of *K*_t_ suggest that environmental conditions in 1997 (the first full year of monitoring) were already sufficient to support a much larger population. Despite this, it took the population at least 15 years to return to its previously observed population size. Similar recovery rates have also been observed in spotted hyenas in the Serengeti (Benhaiem et al., 2018). Population growth between 1997 and 2011 therefore appears to have been a delayed response to environmental improvement that occurred prior to the monitoring period, rather than an immediate response to concurrent environmental change. Our results suggest that conservation actions over the past decades (Table ED1) did not lead to a net improvement in conditions for spotted hyenas in Ngorongoro Crater despite observed population growth. Such conclusions would be impossible to draw using measurements of population size alone.

Our results illustrate that the recovery can occur at different time scales for the environment and for a population. The speed at which a population responds to environmental change varies greatly between species and populations (Ackleh et al., 2018; Coulson and Hudson, 2002; Koons et al., 2007). Slower responses are typical of species with longer generation times that often experience physiological constraints on reproduction (Coulson and Hudson, 2002; Koons et al., 2006, 2007), such as the maximum litter size and the minimum inter-birth interval. The spotted hyena exhibits a late age at first reproduction (3–5 years; supplementary section S1.2.2; Davidian et al., 2016; Gicquel et al., 2022; Holekamp et al., 1996; Höner et al., 2010), long inter-birth interval (20 months average; section S1.2.3), and small litter size (commonly 1–2 cubs Frank, 1986; Holekamp et al., 1996). These features constrain maximum population growth and likely explain the long period of recovery from declines seen in spotted hyenas (Benhaiem et al., 2018; this study). It is not unusual for population sizes of animals or plants to take years or even decades to reflect changes in environmental conditions (Ackleh et al., 2018; Capdevila et al., 2022; Raimondo and Donaldson, 2003). To give another example than spotted hyenas, long-lived red deer on the Isle of Rum took ten years to reach carrying capacity after hunting was banned (Albon et al., 2000; Coulson and Hudson, 2002).

In the most favourable years, we estimate that Ngorongoro Crater could have supported as many as 827 spotted hyenas. However, the population never reached this maximum theoretical size due to periods of poor environmental conditions that caused population declines and led to extended periods of recovery. In agreement with our existing knowledge of the species, we found that infectious disease, prey abundance and competitor density were key factors affecting carrying capacity in spotted hyenas. Infectious disease outbreaks are known to be major sources of mortality in spotted hyenas (Höner et al., 2012; Marescot et al., 2018) and other large carnivores (Kissui and Packer, 2004; Young, 1994). Outbreaks of *Streptococcus* infection in both 2002 and 2019 were major sources of mortality for spotted hyenas in Ngorongoro Crater specifically (Dheer et al., 2022; Höner et al., 2012; Höner et al., 2006). Similarly, variation in prey abundance is a key environmental factor in spotted hyenas (Hofer and East, 1993) and has been proposed as one of the potential causes of spotted hyena decline in Ngorongoro Crater prior to 1996 (Höner et al., 2005). Importantly, conservation efforts since 1996 to limit road use and Maasai pastoralism activity inside Ngorongoro Crater had no discernible effect on carrying capacity, supporting previous findings that pastoralism had no negative effect on spotted hyena recruitment (Dheer et al., 2022) and helping explain the lack of a significant linear trend in *K*_t_ over time.

If prey and competitor population dynamics and the frequency of infectious disease outbreaks remain similar in the future to those of the past 26 years we would expect population size to show similar patterns to those observed since recovery was reached in 2011. However, patterns of prey abundance and infectious disease are likely to shift in the context of climate change, which may impact carrying capacity and thus population size (see section S3.1).

Our case-study of spotted hyenas illustrates the conceptual and practical insights that can be gained by combining *K*_t_ as an integrative measure of environmental conditions with long-term data on population size. *K*_t_ should be thought of as an *enhancement* rather than *replacement* for measuring population size. Large carrying capacity does not guarantee population growth in cases where populations reach such low sizes that their dynamics can be governed by non-environmental factors, such as demographic stochasticity (Lande, 1993) or positive density dependence (i.e., Allee effects; Courchamp et al., 1999; Stephens et al., 1999). Small populations in favourable environmental conditions may still benefit from conservation interventions that directly promote population growth, such as population supplementation (Seddon et al., 2014) or reducing inbreeding depression (Hohenlohe et al., 2021; Kardos et al., 2021). Only by measuring *both* population size and *K*_t_ can appropriate conservation actions be reliably identified.

Individual-based modelling provides a powerful and flexible tool to estimate *K*_t_ that can integrate complex species-specific behavioural and ecological characteristics (e.g., Neil et al., 2020; Ovenden et al., 2019; Piacenza et al., 2017; Radchuk et al., 2021). In theory, other modelling approaches that are capable of population projection—such as matrix-based demographic models—present possible alternative methods to estimate *K*_t_ (see supplementary section S3.2). Regardless of the method chosen, estimating *K*_t_ requires reliable estimates of vital rates. Ideally, vital rate information should be collected under a range of demographic and environmental conditions that approximate as close as possible the conditions that are to be simulated (e.g., varying population size). For long-lived species, the collection of such data necessarily entails the long-term study of individuals in the wild. Such data are difficult and costly to collect (Sheldon et al., 2022) and thus are most likely to be available for large flagship conservation or research programs (e.g., Culina et al., 2021; Morrison et al., 2022). Where possible, missing information may be drawn from other populations of the same species or closely related species with similar life-history characteristics (Coulson and Hudson, 2002; Heppell et al., 2000; Hernández-Camacho et al., 2015).

When collecting individual-level data is infeasible, there is still a possibility to estimate trends in *K*_t_ using ecological proxies such as food availability (e.g., Hayward et al., 2007; Karanth et al., 2004) or territory availability (e.g., Jones et al., 2022). Environmental proxies reduce the need for individual-level data but present their own limitations. Measuring environmental proxies can still impose non-trivial data collection and computational challenges (Jones et al., 2022; Karanth et al., 2004). Unlike *K*_t_, environmental proxies cannot provide information about how close a population is to its demographic equilibrium, which limits any possibility to predict future population growth and abundance. It is also usually unknown how well environmental proxies correlate with *K*_t_. If the correlation between an environmental proxy and *K*_t_ is weaker than expected, relying on environmental proxies can lead to less effective conservation actions. Nevertheless, ecological proxies provide a key alternative method to estimate environmental conditions and supplement information on population size in cases of data limitation.

### 3.1 Conclusions

In this paper, we have proposed time-varying carrying capacity (*K*_t_) as an additional source of conservation information that overcomes potential limitations associated with decisions based on population size alone. *K*_t_ allows for a better assessment of environmental conditions important to a target species or population and provides a more direct metric to assess conservation effectiveness. Our study show how *K*_t_ can be used to disentangle effects of recent environmental change from older environmental change, something which would be impossible using population size alone. Lingering effects of past environmental change are particularly common in long-lived species, making such effects directly relevant for modern conservation, where many charismatic species that garner public attention and investment tend to be long-lived (Clucas et al., 2008; Davies et al., 2018). In sum, quantifying *K*_t_ expands the toolkit available to conservation managers and facilitates evidence-based conservation outcomes, which is vital to most effectively address the global biodiversity crisis.

## 4 **Methods**

### 4.1 Study species and population

The spotted hyena (*Crocuta crocuta*) is a large social carnivore that plays an important role in the African savannah ecosystem as a regulator of herbivore disease (Packer et al., 2003), a competitor of other large carnivores (Höner et al., 2002), and as a species capable of digesting large bones and thus releasing calcium and phosphorus concentrated within bones back into the ecosystem (Abraham et al., 2022). Although the spotted hyena is categorised as a species of ‘Least Concern’, many populations are declining due to impacts of human activity such as culling, poisoning, snares, and roadkill (Benhaiem et al., 2023; Bohm and Höner, 2015; Naciri et al., 2023). Spotted hyenas form clans, consisting of multiple matrilines organized into a relatively stable linear dominance hierarchy (Kruuk, 1972), where cubs obtain a social rank within the dominance hierarchy directly below that of their mother and above older siblings (Frank, 1986; Vullioud et al., 2019). Dispersal is strongly male biased (East and Hofer, 2001; Höner et al., 2005). Most males disperse to new clans once reaching adulthood, where they join the bottom of the social hierarchy (Höner et al., 2007), although male philopatry is also observed (Davidian et al., 2016).

Reproductive success in spotted hyena females is predominantly determined by the position of an individual in the social hierarchy (e.g., Frank, 1986; Höner et al., 2010)—as evidenced by the large skew observed in reproductive success between high and low ranking females (Holekamp et al., 1996). Females reproduce year round, with a gestation time of around 110 days (Matthews, 1939). Litter size ranges from one to three cubs, although competition between siblings means that subordinate siblings in multi-cub litters have a reduced survival (Gicquel et al., 2022; Hofer and East, 2008). Cubs are typically not weaned until 12–20 months of age (Hofer and East, 1995; Holekamp et al., 1996; Watts et al., 2009), and are generally considered sexually mature around 24 months (Matthews, 1939) although age at first reproduction usually occurs between 3–5 years old (section S1.2.2; Davidian et al., 2016; Gicquel et al., 2022; Holekamp et al., 1996; Höner et al., 2010).

The population of spotted hyenas in the Ngorongoro Crater, Tanzania (3°12’36”S, 35°27’36”E) includes eight clans that defend territories within the 250km^2^ area of Ngorongoro crater floor. The population has been monitored since April 1996 as part of the Ngorongoro Hyena Project (https://hyena-project.com). Individuals are identified by their unique spotting patterns as well as distinct characteristics like ear notches and scars. Observations of dyadic interactions and genetic-based parentage analyses have allowed for an accurate estimation of the social hierarchy within each clan (Vullioud et al., 2019). Extensive behavioural observations have provided a strong understanding of both male dispersal and female mate choice behaviour (Davidian and Höner, 2022; Davidian et al., 2016; Höner et al., 2007).

### 4.2 Estimation of population size and its trends over time

To estimate population size (*N*_t_), we counted the total number of spotted hyenas known to have been a resident of at least one of the eight clans inhabiting the floor of the Ngorongoro Crater during a specific time period. Note that a given hyena may have belonged to more than one clan if dispersal occurred during that period. Unless mentioned otherwise, *N*_t_ values refer to annual estimates, which means that they include all individuals that have been resident for a least one day during year t. Individuals are considered as resident either when they are born in the clan and have not yet dispersed or died, or have immigrated into a clan. An individual is considered to have immigrated when they are observed, during at least three months, involved in affiliative social interactions with members of that clan or otherwise observed at the communal den.

Birth, death and dispersal events are rarely directly observed in the wild. These events have thus been estimated and cross-checked using multiple proxies derived from raw data collected during near-daily visits. These data have been collected by five experienced field observers. Proxies of life history events include: (re)sightings; morphological and behavioural observations; genetic parentage analyses of biological samples (for details, see Davidian et al., 2016; Dheer et al., 2022). A few transient individuals, originating from unknown clans or known clans outside the Crater, and not known to be socially integrated in any Crater clans, are not considered members of the population and are thus not included in *N*_t_ estimates.

To quantify temporal trends in population size, we modelled change in *N*_t_ over time with a linear model including a fixed quantitative year term. Due to the potential for temporal auto-correlation of residuals when modelling *N*_t_, we also included an AR1 term to account for correlation between successive time steps. We tested separately for trends in *N*_t_ across the total study period (1997–2022), across the period of initial population growth (1997–2011), and across the subsequent period (2012–2022). These models assumed a Gaussian distribution of residuals with mean zero and variance *σ*^2^.

All analyses of linear models and Generalised Linear Mixed-effects Models (GLMMs) present in this paper were done using the R package spaMM (Rousset and Ferdy, 2014). We performed all tests of fixed effects in such statistical models using asymptotic log-likelihood ratio tests and calculated the 95% confidence interval (CI_95%_) using likelihood profiling.

### 4.3 Spotted Hyena Individual-based Model (SHIM)

Individual-based models (IBMs) are suitable for modelling complex systems and are used across a range of fields (Farmer and Foley, 2009), including in conservation science (e.g., Boult et al., 2018, 2019; Neil et al., 2020; Ovenden et al., 2019; Piacenza et al., 2017). IBMs integrate effects of demographic and environmental stochasticity. They can also integrate a wide range of mechanisms that may influence population dynamics, including complex animal life-cycles (Piacenza et al., 2017) and extrinsic pressures such as climatic change (Boult et al., 2019).

To study time-varying carrying capacity (*K*_t_) in our study population, we created an IBM that simulates the eight spotted hyena clans within the Ngorongoro Crater. We named this model the ‘Spotted Hyena Individual-based Model’ (SHIM). We present a brief overview of the simulation model here, but a full description of the model following the ‘Overview, Design Concepts, and Details’ (ODD) framework (Grimm and Railsback, 2012; Grimm et al., 2006) can be found in section S1.1.

SHIM simulates the life cycle and behaviour of spotted hyenas using discrete time steps (months). In a given month, individuals may reproduce (give birth), die, and disperse—whichever are appropriate for their particular age and sex. These events are determined by stochastic predictions stemming from seven GLMMs fitted on observed data from Ngorongoro Crater collected since 1996. These statistical models estimate effects of individual characteristics such as age, sex, and rank within the hierarchy, on the survival, reproduction, twinning, and dispersal of spotted hyenas (for details, see section S1.1.7; Table ED2).

In each time step, sexually mature females (*≥* 24 months) attempt to select a mate following typical mate choice patterns documented in this population (Höner et al., 2007). If at least one suitable male is present in the clan, females have a chance to reproduce with a probability determined by one of the two statistical models for reproduction—one for primiparous females and one for non-primiparous females. Suitable males must be adults (*≥* 24 months) and have been born or immigrated into the clan after the female was born. Females that reproduce may have a litter with one or two cubs. Cases of three-cub litters are rare in the wild and not considered within the simulation. Each new cub from a reproduction event is given a rank just below that of their mother and above their older siblings following the social inheritance behaviour documented in the system (Frank, 1986).

Field observation of cubs in their first few weeks of life are difficult as they spend most of their time within their birthing den. Young cubs may thus die before ever being observed. To circumvent limitations caused by the unreliable estimation of cub survival, we chose to integrate early life survival into our measure of reproduction. In our reproduction models, we therefore defined a reproduction event as a female that gives birth to a litter where at least one cub survives to six months of age. In the simulation, this means that all cubs from a successful reproduction event are assumed to survive for at least six months.

The probability that a female has a multi-cub litter is determined by our statistical model for twinning. This statistical model also integrates early life survival such that a multi-cub litter is a litter where more than one cub survives to at least six months of age. In our statistical model, we grouped observed litters of both two and three cubs together, which allowed us to work with a binomial response variable. As mentioned above, cases of three cub litters are rare in the wild and so all multi-cub litters in the simulation have a litter size of two. For convenience, we refer to the probability of multi-cub litters as ‘twinning’ throughout the manuscript, although multi-cub litters with mixed paternity are possible.

All individuals six months or older may die during a time step with a probability determined by our statistical models for survival. Cubs younger than six months are excluded from this process as early life survival is already integrated into our statistical models for reproduction and twinning. We used separate survival models for females, juvenile males (*<* 24 months), and adult males (*≥* 24 months). Due to the structure of our male dispersal model (described below) all adult males are necessarily males that have selected a clan (including philopatric males).

Males reaching 24 months are assumed to be adults. At this point, they prospect for a possible new clan in which to disperse. The clan a male chooses is determined by a multinomial model, where the probability of selecting a clan is a function of the relative number of females between one and five years old present in the clan (Höner et al., 2007). The male’s birth clan is considered as a possible choice, allowing for the possibility of philopatry (Davidian et al., 2016). If a male enters a new clan (i.e., a non-philopatric male), he is placed at the bottom of the social hierarchy. In contrast, philopatric males keep their previous social rank upon reaching 24 months.

Once a male has selected his first clan (either philopatric or non-philopatric), he has a probability to disperse again at each subsequent time step determined by our statistical model for additional male dispersal. In the field, it is difficult to identify cases where adult males might prospect new clans but do not ultimately disperse. Therefore, this statistical model only considers cases where males move from their current clan to a new clan, and so males cannot reselect their current clan. The probability with which a male selects a new clan is again drawn from a multinomial model which is a function of the relative number of young females (between one and five years old) in all clans but their current one. Males that disperse from their birth clan and return to their birth clan after having lived elsewhere are not considered philopatric (i.e., they join the bottom of the social hierarchy after dispersal back into their birth clan).

We undertook a pattern oriented modelling approach (Grimm and Railsback, 2005, 2012), to demonstrate the reliability and biological realism of SHIM. We compared the emergent properties of our model to observed demographic patterns in the wild, including patterns of lifetime reproductive success and lifespan of both sexes. SHIM was able to recreate a range of complex demographic patterns observed in Ngorongoro Crater. These comparisons are detailed in section S1.2.

### 4.4 Estimation of the time-varying carrying capacity

We used SHIM to estimate *K*_t_ in Ngorongoro Crater. All statistical models, except for that of male additional dispersal, included inter-annual variability in vital rates through the inclusion of a random effect when simulating life history events using the fitted statistical models. To estimate *K*_t_ for a given year (t), we ran SHIM with environmental conditions kept constant to those observed in year t by fixing the inferred realisation of the random effect for that year. In each simulation run, SHIM continued for 1,200 time steps (100 years). Stochasticity in demographic outcomes due to individual characteristics (e.g., age, rank) was included. The value of *K*_t_ for a single simulation run was computed as the median population size once the population was considered to have reached equilibrium. We used this method to estimate carrying capacity for every year from 1997 to 2022. The year 1996 was excluded as the population was not monitored for the full year. We also excluded the most recent year (2023) as we considered estimates of survival and reproduction to be too uncertain for that year due to fewer opportunities for resightings and time required to process genetic samples used for parentage assignment.

For each year, we carried out ten independent simulation runs. Each simulation was initialized using the same snapshot of the population observed in April, 1996 (178 individuals), including correct age, sex, rank, and parental relationships. Our final estimate of *K*_t_ in each year was then the median value of *K*_t_ across all ten replicates. In this way we could estimate *K*_t_ across the study period.

To remove initialization bias (i.e., impact of starting conditions) the first half of each simulation run (50 years) was considered a burn-in period and discarded before estimating *K*_t_. Following the removal of the burn-in period, visual inspection of population size over time was carried out for all replicates to check for trends in population size that would suggest carrying capacity had not been reached. We considered carrying capacity to have been reached in all simulation runs following visual inspection and no further action was taken.

### 4.5 Comparison to estimates assuming a fixed carrying capacity

We compared our estimate of *K*_t_ from SHIM with methods that assumed a fixed carrying capacity over time. We reran SHIM using the same statistical models as described above but where stochastic outcomes were produced by marginal rather than by conditional simulations. To produce marginal simulations, we first computed marginal predictions of the vital rates, which are functions of coefficients of the fixed effects, and of the variance of the random effects. Specifically, we averaged, over the fitted distribution of random effects, the predictions of our vital rates expressed on the scale of the response (i.e., back-transformed from the scale of the linear predictor) and conditional on the fixed and random effects. Unlike the conditional predictions computed for a specific value of the random effects (typically zero), such computation provides unbiased predictions. It should thus be favoured in the context of GLMMs where random effects act non-additively on the expected response (Bland and Cook, 2019). To turn these marginal predictions into marginal simulations of the vital rates, we then drew random binary events using the function *rbinom()* from the R package stats using the marginal predictions as inputs for the probabilities. We used the same starting population, number of simulation runs, and number of time steps for our simulation as described above.

We also estimated fixed carrying capacity using the traditional Ricker (Ricker, 1954) and Beverton-Holt (Beverton and Holt, 1957) demographic models. We considered both the exponential and hyperbolic version of these models (Johst et al., 2008) (see section S2.3 for details).

### 4.6 Temporal trends in carrying capacity

To quantify temporal trends in carrying capacity, we modelled change in *K*_t_ over time similarly to how we modelled change in *N*_t_, i.e., with a linear model including a fixed quantitative year term. We also modelled temporal trends in carrying capacity at the clan level (*K*_tc_) using a linear model that included a quantitative term for year interacting with clan. As for the study of *N*_t_, we also included an AR1 term to account for correlation between successive time steps when modelling *K*_t_ or *K*_tc_. We also tested for trends in both *K*_t_ and *K*_tc_ during the period of initial population growth (1997–2011).

### 4.7 Time-varying carrying capacity and environmental drivers

To assess how changes in the environment could explain variation in *K*_t_ we fitted a statistical model with carrying capacity at the clan level (*K*_tc_) as the response variable and five potential candidate predictors. Predictors were selected *a priori* so as to capture major potential drivers of population change (inter-specific interactions, disease and anthropogenic impacts/conservation measures). We conducted analysis at the clan level so that we could account for variation in predictors expected between clans (e.g., heterogeneity in abundance of prey and competitors).

We included a semi-quantitative estimation of prey abundance based on the ordinal ranking of prey abundance around spotted hyena den sites, recorded during every den visit (1 = low density; 2 = medium density; 3 = high density). To estimate prey density for a clan we treated these categories as numeric variables and took the average across all den visits in a year. To test for potential effects of competitors, we included a quantitative estimation of the abundance of lions in each clan and year. We first estimated territory boundaries of each spotted hyena clan using 95% minimum convex polygons applied to sightings of adult females. Adult females were used because sightings of males may include males prospecting or dispersing to new clans, which is not indicative of clan space use. Clan territories were calculated separately for two periods, 1996–2011 and 2012–present. These different periods were used to account for known shifts in prey space use within Ngorongoro Crater, which we expected would also affect spotted hyena territories. More details about shifts in prey and justification for using different periods are provided in Dheer et al. (2022). Once territories were calculated, we counted the total number of lions sighted within each clan territory in each year and determined the lion density (lions/km^2^) within each territory. Prevalence of infectious disease was quantified in each clan each year using sightings of spotted hyenas with clinical signs of infectious disease. For example, *Streptococcus* infection is associated with clinical signs that can be clearly identified in the field (Höner et al., 2006). In each year, we calculated the proportion of individuals in each clan observed with clinical signs of disease. To account for effects of conservation management, we included two binary variables that account for whether certain conservation measures were in effect during a year (whether or not tourist road use was restricted and whether or not Maasai pastoralists were allowed to visit the Crater with cattle; Table ED1).

To test the potential drivers of carrying capacity we fitted a linear mixed-effects model. We included all five variables described above (lion density, prey abundance, disease prevalence, road use restrictions, pastoralism ban) plus a categorical term for clan to account for differences in quality between clan territories. We included an AR1 term to account for temporal auto-correlation between successive time steps.

## 5 Code availability

All code used for analyses is available through the GitHub repository Bailey2024K (https://github.com/hyenaproject/Bailey2024K).

## 6 **Data availability**

All data required to reproduce analyses has been deposited on Zenodo (https://zenodo.org/ doi/10.5281/zenodo.10955614).

## 7 Author contributions

**CRediT (Contributor Roles Taxonomy) author statement.** Note the “Project” is here considered to refer to the current study. ED also contributed to the project administration and funding acquisition of the larger Ngorongoro Hyena Project.

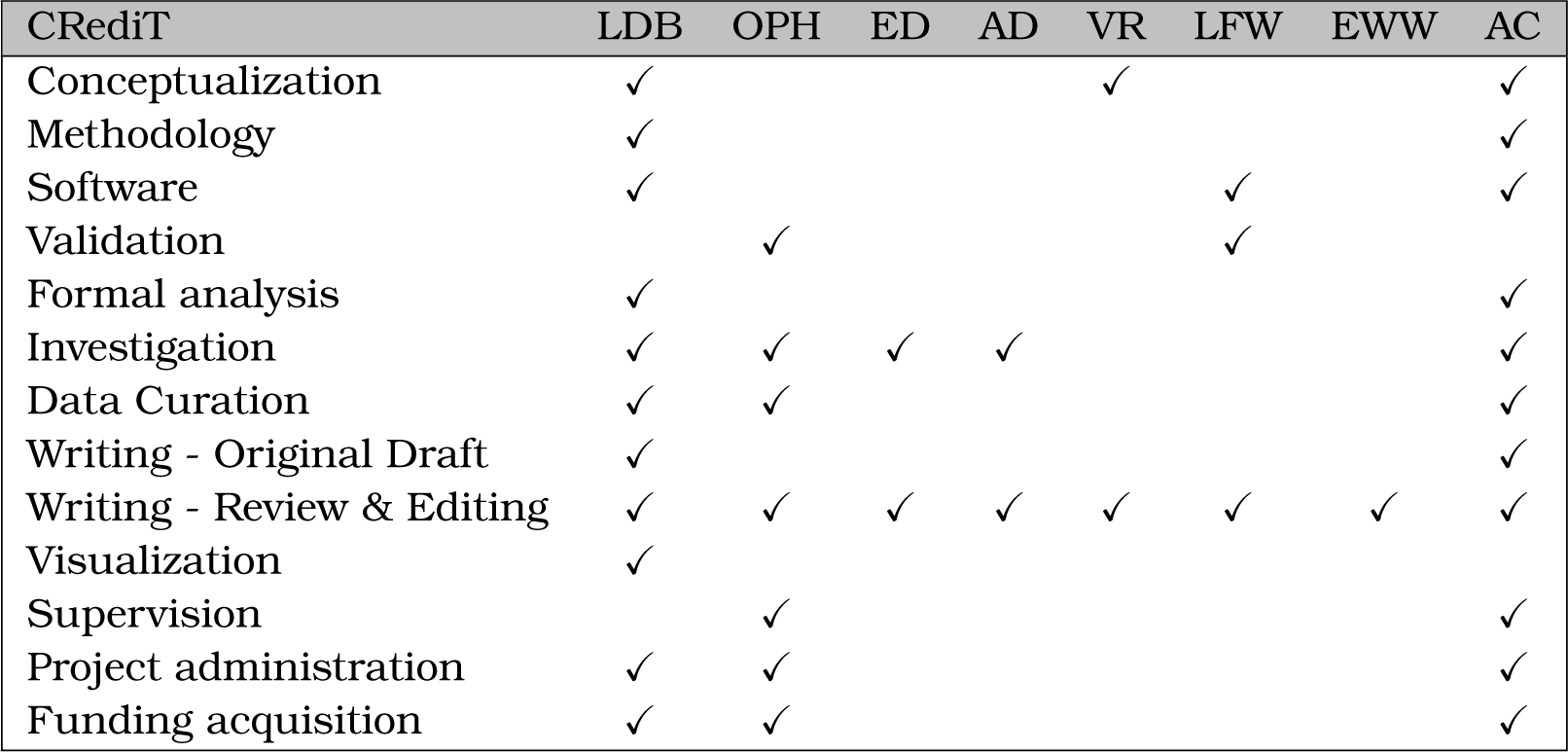

## Supporting information

Supplementary Methods, Results and Discussion

## Acknowledgements

Many thanks to Sarah Benhaiem, Olivia Judson and Franç ois Rousset for feedback on the manuscript. Additional thanks to Franç ois Rousset for advice on statistics as well as the development of the R package spaMM. Thanks also to Philemon Naman and Bettina Wachter for their invaluable contributions to data collection and project management. We thank the Tanzania Commission for Science and Technology, the Tanzania Wildlife Research Institute and the Ngorongoro Conservation Area Authority for permission to conduct the study. This study was funded by the Deutsche Forschungsgemeinschaft (DFG, German Research Foundation)— BA 6703/2, and the Leibniz Institute for Zoo and Wildlife Research.

## Extended Data

**Table ED1:** Important events in Ngorongoro Crater, Tanzania. Rows below the horizontal line show events that have occurred during the study period.

| Year | Event | Reference |
| --- | --- | --- |
| 1959 | NCA established as multiple land-use area | UNESCO World Heritage Convention |
| 1964–1968 | First study of spotted hyenas in Ngorongoro Crater | Kruuk (1972) |
| 1974 | Maasai no longer allowed to stay over night in Ngorongoro Crater |  |
| 1979 | NCA inscribed as UNESCO World Heritage | UNESCO World Heritage Convention |
| 1980s | Beginning of mass tourism |  |
| 1980s–90s | Culling of spotted hyenas | Personal communication |
| 1996 | Ngorongoro Hyena Project established | Höner et al. (2005) |
| 1998 | Heavy El Niño rains | Estes et al. (2006) |
| 2000 | Severe drought | Kissui and Packer (2004) |
| 2001 | Canine distemper virus (CDV) outbreak among Ngorongoro Crater lions | Kissui and Packer (2004) |
| 2002–2003 | *Streptococcus* outbreak among spotted hyenas | Höner et al. (2006) |
| 2006 | Restriction on road use | Personal observation |
| 2007–2008 | Eruption of volcano Oldoinyo Lengai | De Schutter et al. (2015) |
| 2016 | Maasai no longer allowed to bring livestock into Ngorongoro Crater | Melubo and Lovelock (2019)<br>Dheer et al. (2022) |
| 2019 | *Streptococcus* outbreak among spotted hyenas | Personal observation |

**Table ED2:**
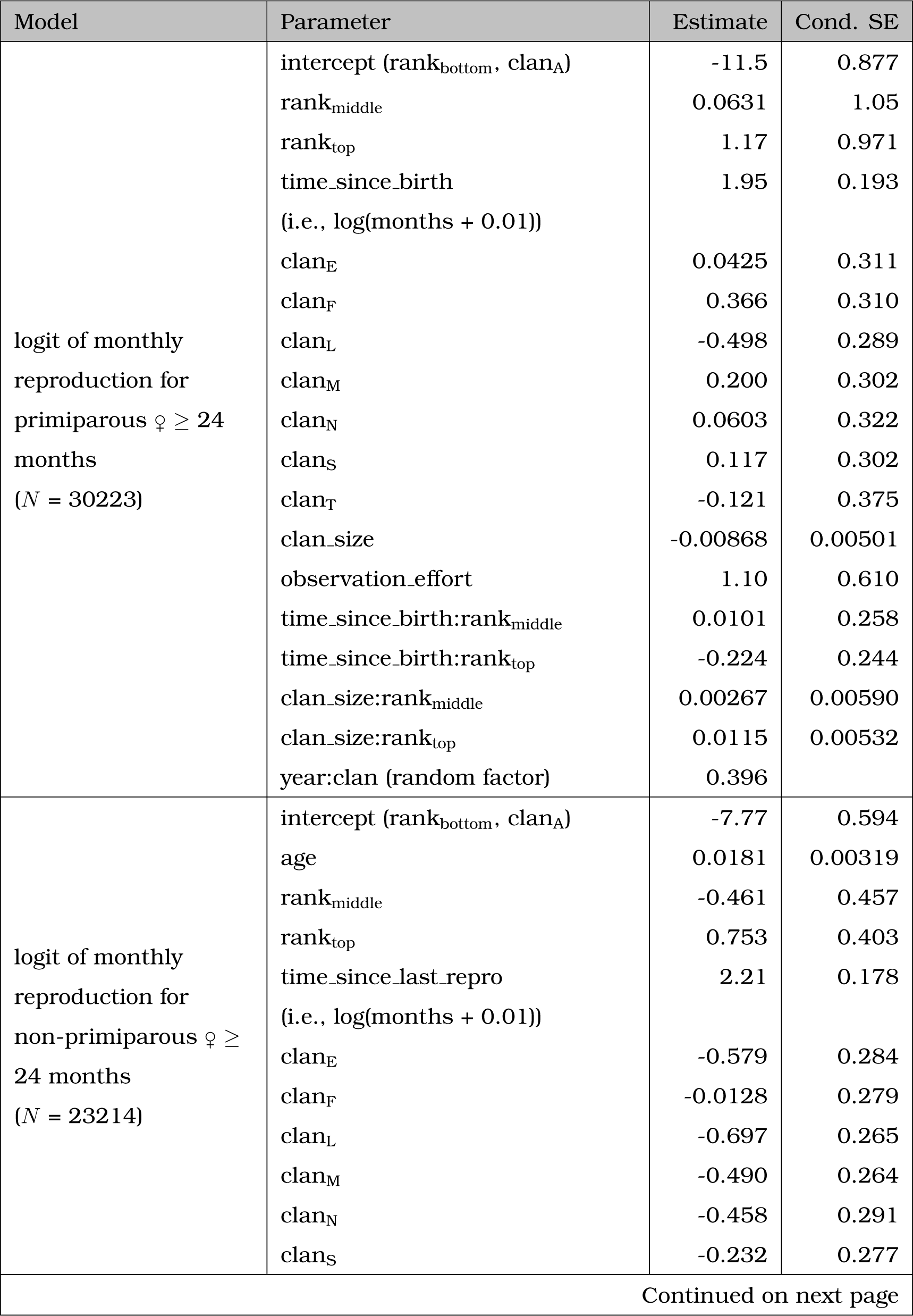
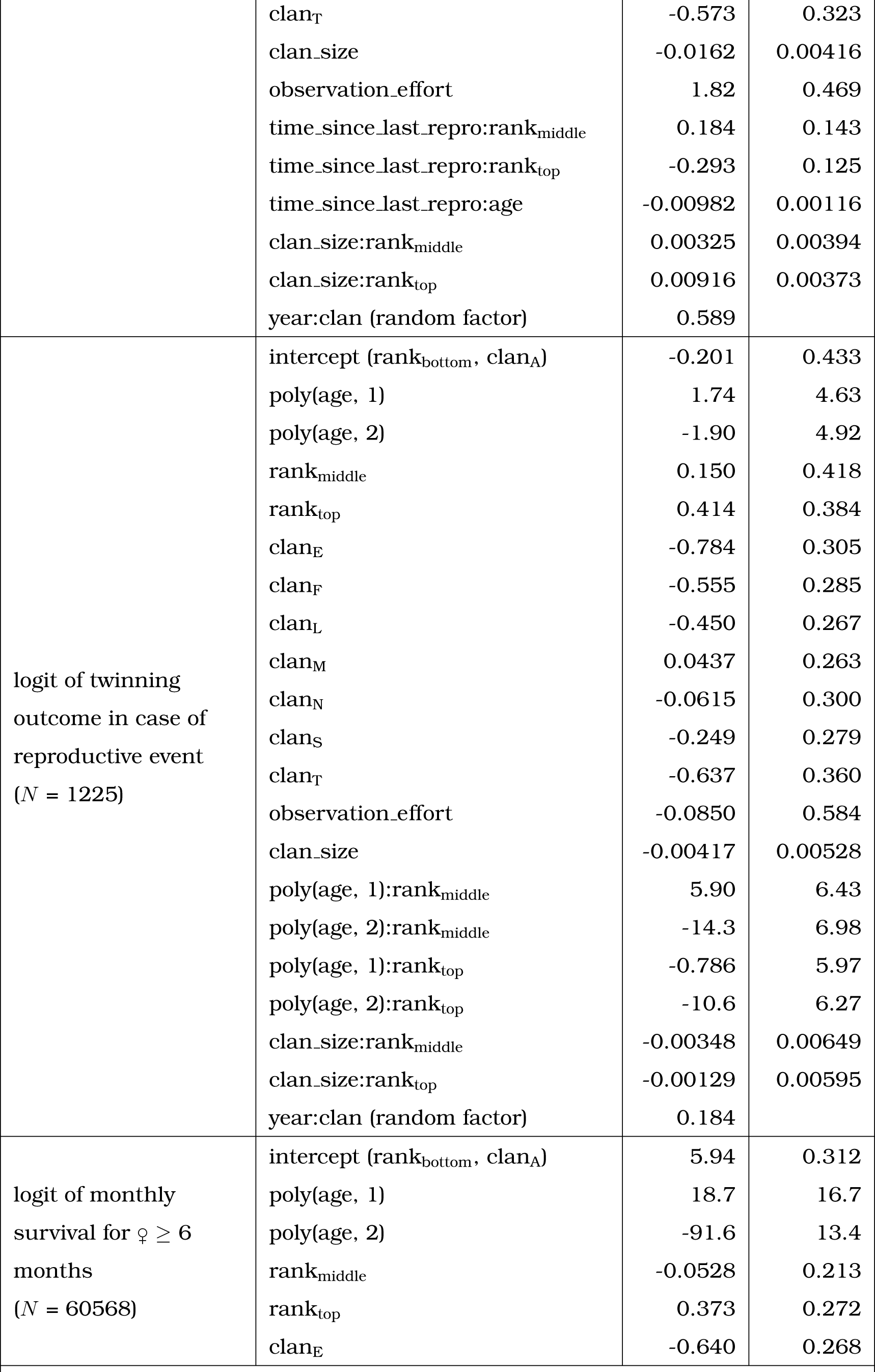
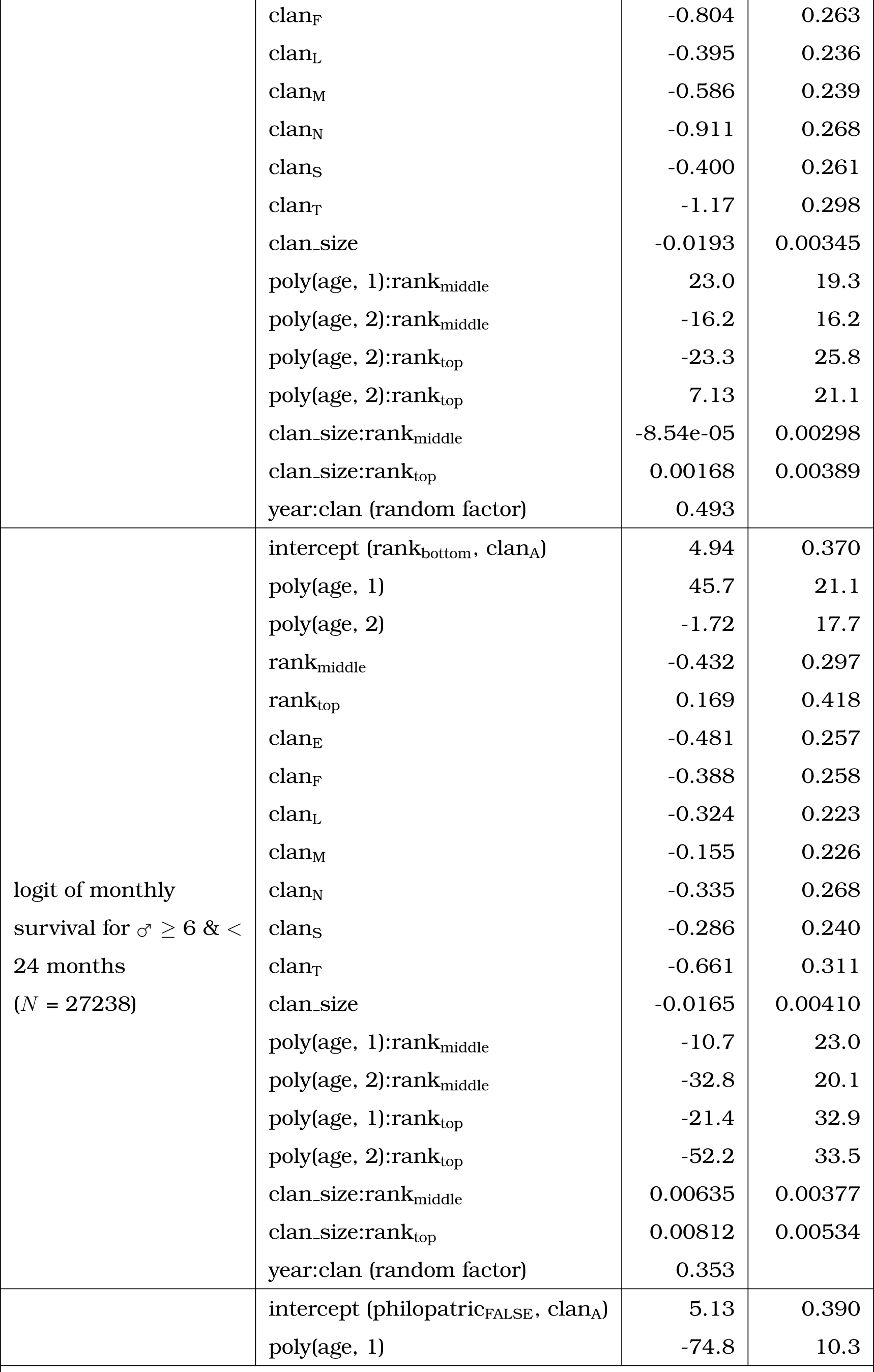
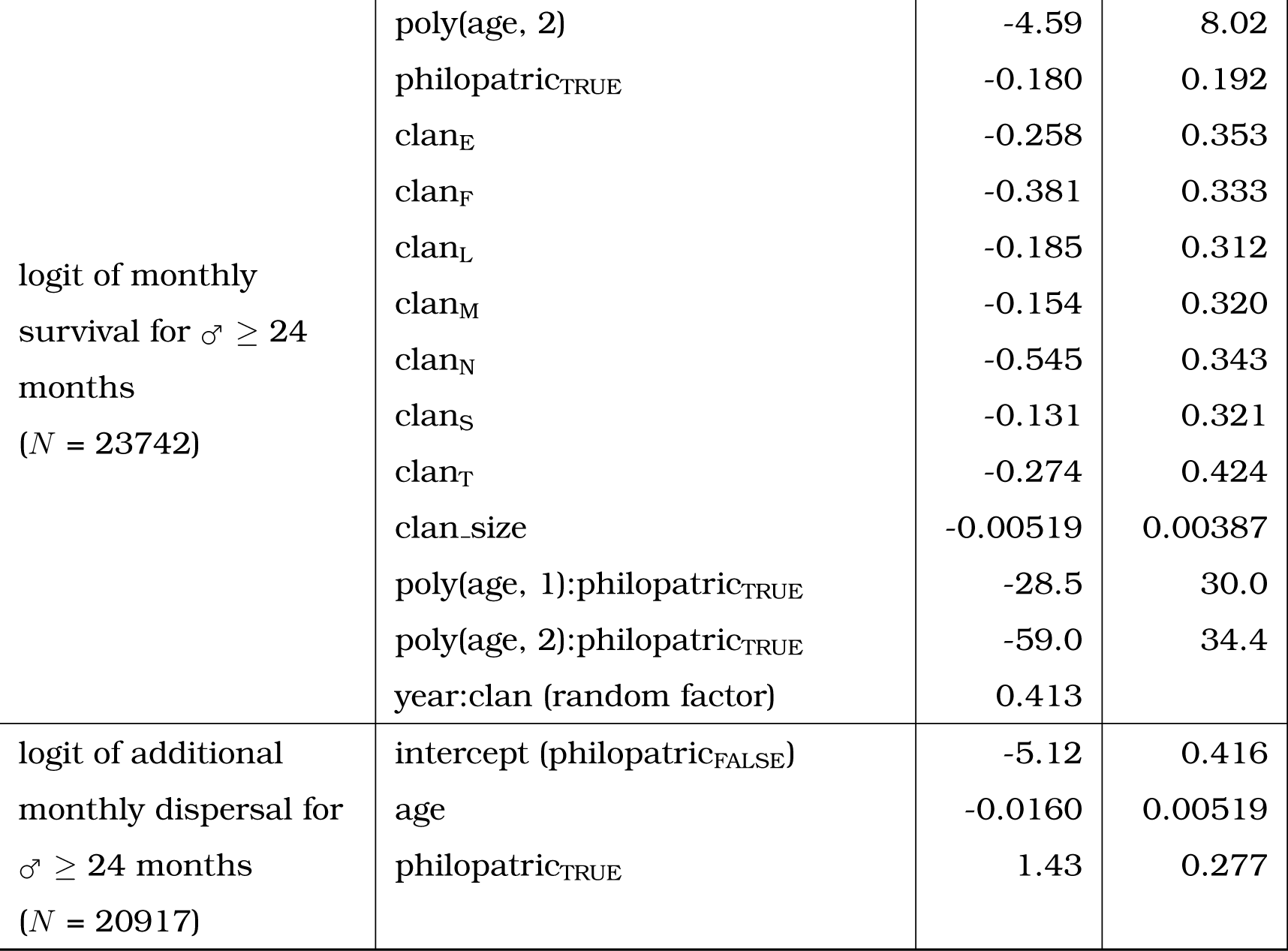
Estimates of vital rates models. This table provides, for each life history trait modeled, the estimate values for each parameters and their associated conditional standard errors. For **models** including both ‘poly(age, 1)’ and ‘poly(age, 2)’, estimates for these parameters are given for orthogonal polynomials produced by the function *poly()* in R and thus do not correspond to raw polynomial estimates. For ‘year:clan’ the variance of the random effect is provided as an estimate and no conditional standard error is given. All fitted events correspond to binary events. All sample sizes (*N*) refer to the number of such fitted events.

**Figure ED1:**
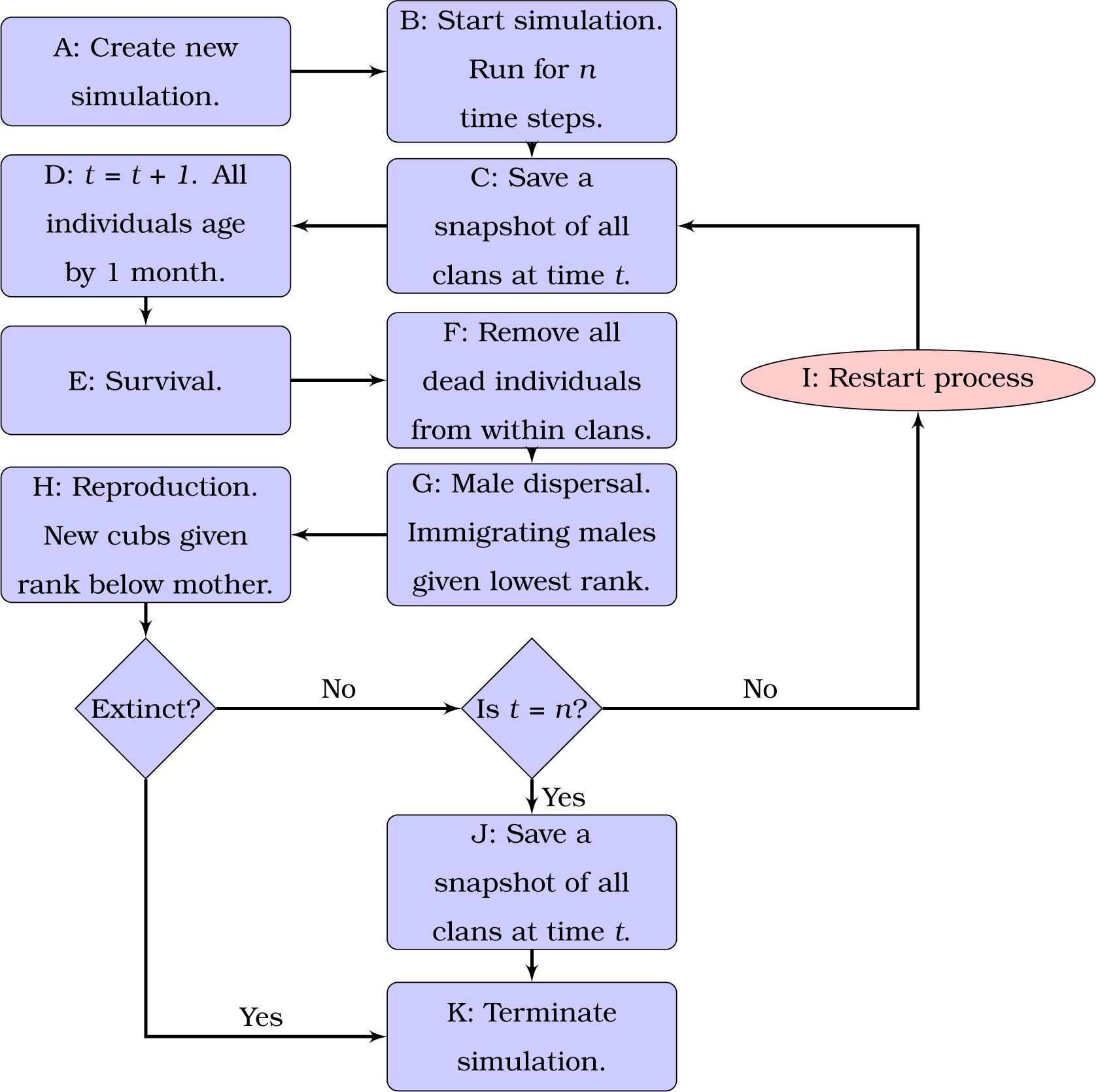
Flow chart of the Ngorongoro Crater Spotted Hyena Individual-based model (SHIM). (A) Simulation is created and initialization conditions are defined. (B) Simulation begins. Simulation is run for *n* time steps or until the simulated population is extinct. (C) A snapshot of individuals in all clans is saved. (D) Time is advanced by 1 month. (E) Survival is determined for all individuals. Probability of survival is predicted separately for females, juvenile males (*<* 24 months), and adult males (*≥* 24 months). (F) Dead individuals are removed. These individuals do not count towards density dependent effects in statistical models of vital rates. (G) Adult males disperse based on availability of young females. (H) Females reproduce with a probability determined by female reproduction models. Females select a male based on observed mate choice patterns. Litter size can be either 1 or 2, determined by twinning model. (I) If the population is not extinct and number of time steps (*n*) has not been reached the process restarts. (J) If number of time steps (*n*) has been reached, a final snapshot of the population is saved. (K) Simulation is terminated.

**Figure ED2:**
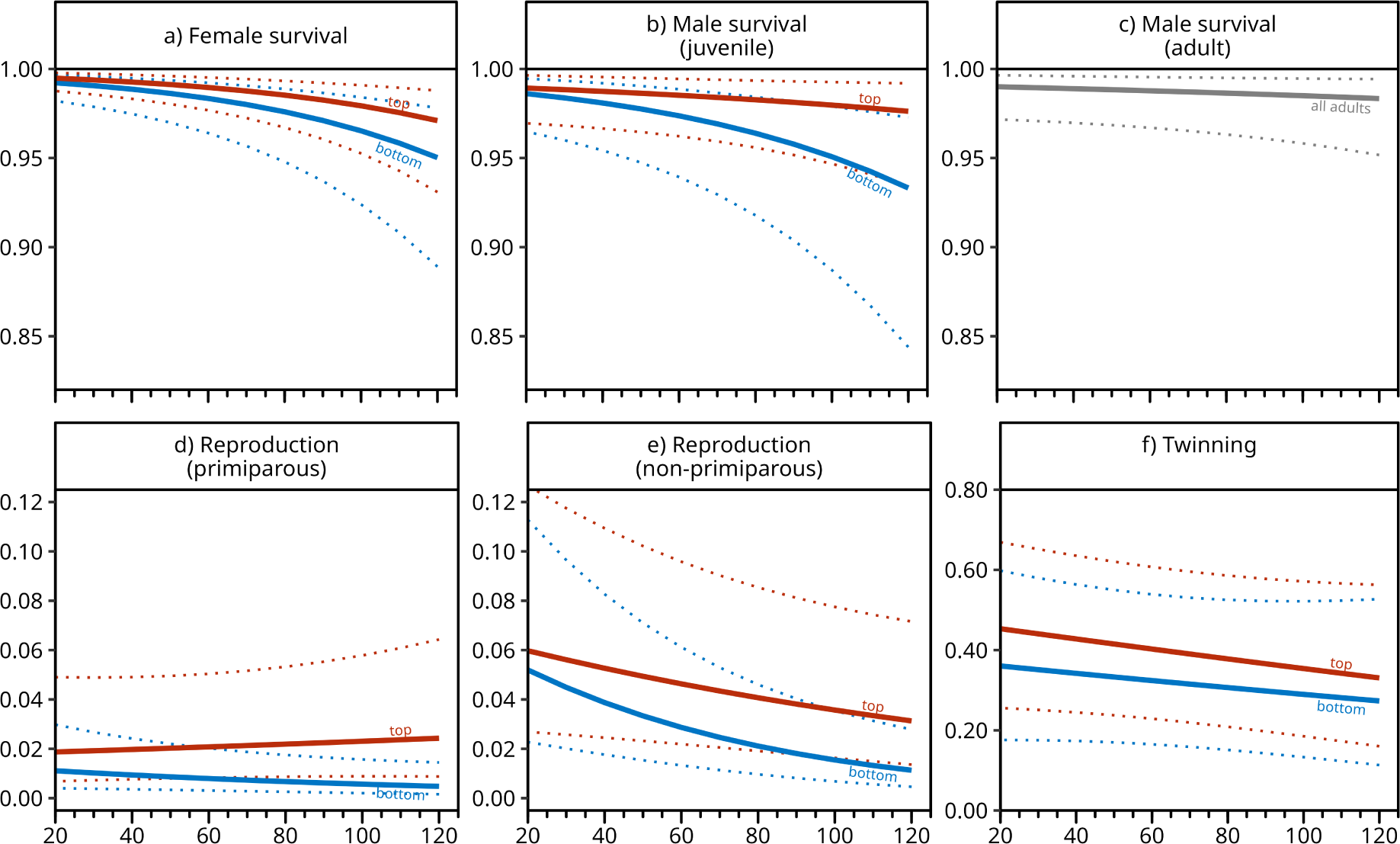
Density dependence in vital rate models. Solid lines depict the predicted probability of life history events shaping the survival and reproduction of spotted hyenas differing in social ranks between bottom ranking individual (blue) and top ranking individuals (red). The x-axis represents the number of individuals in a clan. The dotted lines show confidence intervals (95%) around the predictions. All predictions were computed, for each value sampled along the x-axis and for both top and bottom ranking individuals, as the average of predicted values on the response scale, over the empirical distribution of all other fixed-effect variables and of inferred random effects (i.e., as partial dependence effects as defined in the R package spaMM). This method allows for the visualisation of the effect of specific predictor variables while avoiding any effects caused by association with other predictor variables. Note that the vital model predicting the survival of adult males (f) did not consider distinct categories of social rank since most would be classified as bottom ranking. Note also that the range of the y-axis is not the same for all panels. See Fig. S7 and Fig. S8; for additional results on the relationship between social rank and density dependence, including middle ranking individuals.

**Figure ED3:**
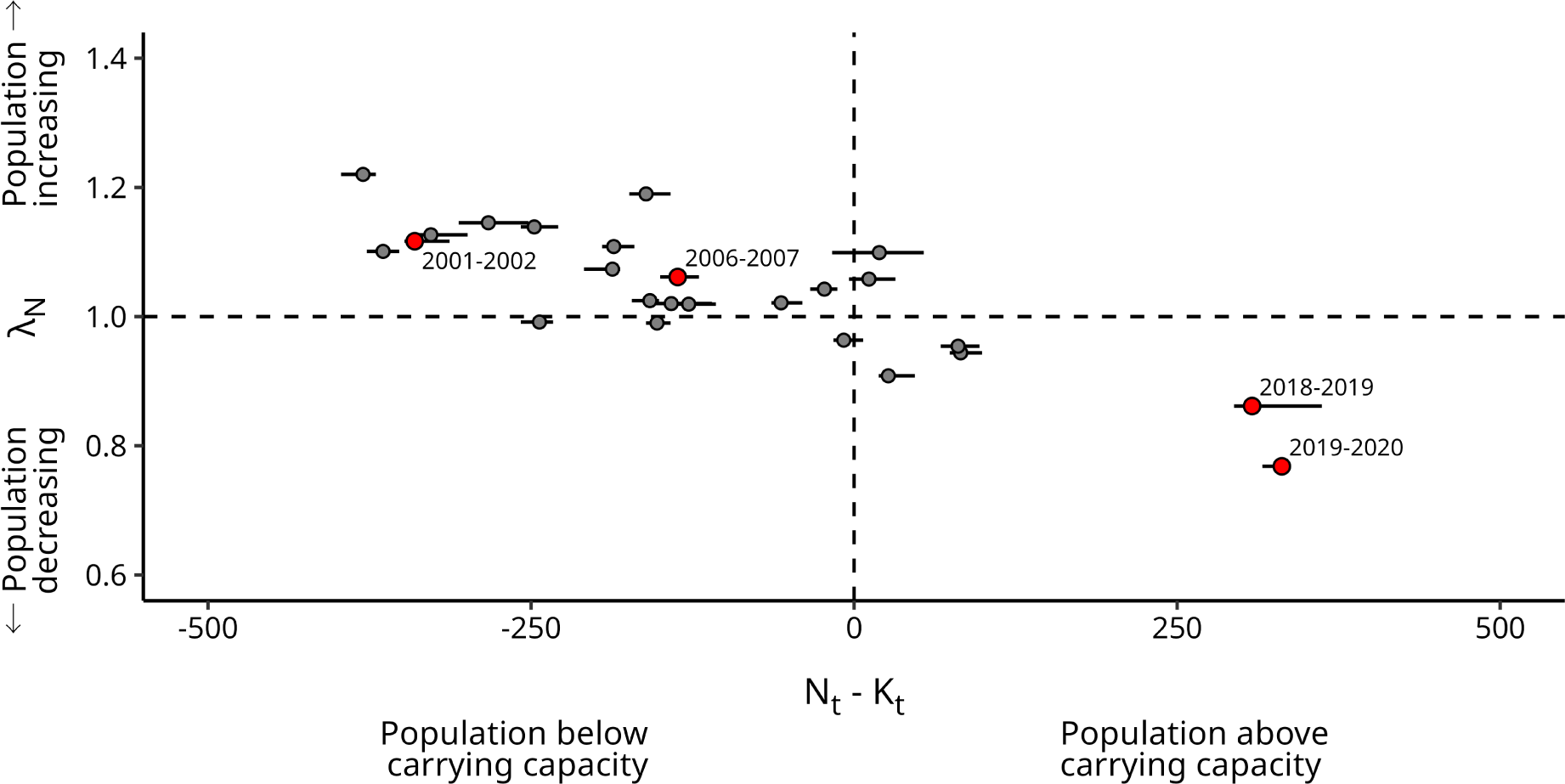
Inter-annual change in population size (λ*_N_*) is strongly correlated with *N*_t_ *− K*_t_. Relationship between annual population growth (*λ_N_*) and the difference between carrying capacity and population size at the start of the time period. See Fig. S11b for plot with year labels for all points.

**Figure ED4:**
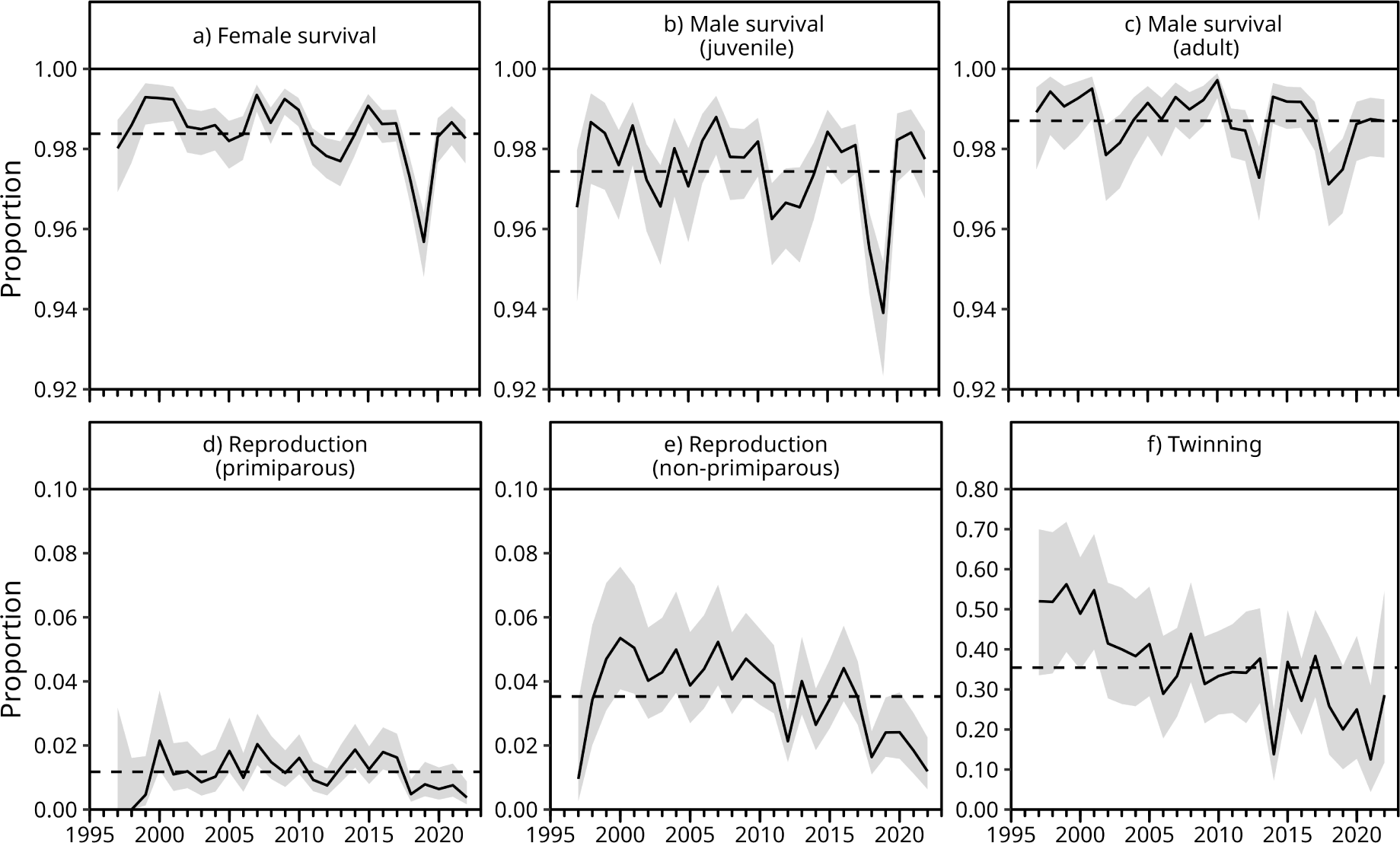
Vital rates of spotted hyenas show no positive linear trends over the years. The solid lines represent the observed monthly proportion of ‘successful’ outcomes for each vital rate, with shaded regions showing 95% confidence intervals calculated using the Wilson method (Wilson, 1927). Dashed horizontal line represents the mean of each vital rate over the full study period. We observed no significant linear change in survival or twinning over time. Rates of reproduction significantly declined when accounting for additional co-variates (e.g., age, clan) as shown in partial dependence plots in Fig. S12. Note that the range of the y-axis is not the same for all panels.

