## Supplementary Methods, Results and Discussion for "Effects of environmental change on population growth: monitoring time-varying carrying capacity in free-ranging spotted hyenas"

April 30, 2024

### S1 Supplementary Material 1: Additional methods

#### S1.1 Overview, Design concepts and Details (ODD)

Below we describe the ‘Spotted Hyena Individual-based Model’ (SHIM) using the ODD framework (Grimm and Railsback, 2012; Grimm et al., 2006). This is intended to provide a detailed overview of SHIM. A more general description is included in Methods.

##### S1.1.1 Purpose

The purpose of SHIM is, in this study, to estimate time-varying carrying capacity ($K_t$) of spotted hyenas in Ngorongoro Crater, Tanzania from 1997 to 2022 at a yearly resolution.

##### S1.1.2 State variables and scales

The model compromised three hierarchical levels: individuals, clans, and the population.

Individuals have the state variables: ‘ID’; ‘sex’; ‘age’ (months); ‘current clan’; ‘birth clan’; ‘motherID’ and ‘fatherID’; ‘tenure’ (number of months spent in current clan); and ‘social rank’. Males and females below 24 months are classified as juveniles. All other individuals are considered adults. Adult males ($\geq$ 24-months old) prospect for new clans and have the additional state variable ‘post-dispersal status’, which can be either philopatric (adult males remaining in their birth clan) or disperser (adult males that have dispersed from their birth clan). Adult females are considered reproductively active and have the additional state variables ‘first reproduction’ (date of first birth) and ‘last reproduction’ (date of most recent birth). As juvenile females ($\leq$ 24 months) are not considered reproductively active they have a probability of reproduction fixed at 0 (Table S1). A female is considered to have reproduced if she produces a litter where at least one cub survives to six months old, thus incorporating both probability of reproduction and early life survival. Monthly survival of females, juvenile males, and adult males is determined by separate survival models (see section S1.1.7). Survival models never apply to cubs younger than six months as this is incorporated into the female reproduction model. Out of all the individual state variables, the information provided by ‘sex’, ‘age’, ‘current clan’, ‘social rank’, ‘post-dispersal status’ and ‘last reproduction’ are used in the vital rate models detailed below (see section S1.1.7).

Clans include the state variables: ‘clan’, ‘clan size’, and a list of individuals present. The variable ‘clan’ provides the name of the clan among the eight real clans observed in Ngorongoro Crater and is used as a predictor variable in all vital rate models (see below). The variable ‘clan size’ is used as a predictor variable in vital rate models to incorporate density dependent effects. If clan size reaches 0, a clan is considered to be extinct and cannot be re-colonized. Individuals in each clan are ordered along a strict linear dominance hierarchy. New cubs are placed just below their mother, with younger cubs given a higher rank than their older siblings. Immigrant males are placed at the bottom of the social hierarchy.

The highest hierarchical level in the model is the population. The population has two state variables: a list of the eight clans and the date. Clans are not spatially explicit within SHIM, and male spotted hyenas can disperse to any other clan. The clan a male chooses is affected by reproductive opportunities available in a given clan (see section S1.1.4). For simplicity, the population is treated as a closed system, which excludes the possibility of immigration and emigration of spotted hyenas outside of Ngorongoro Crater.

Table S1: **Overview of parameters and default values.** Dimensionless parameters are either counts or probabilities.

| Parameter | Value |
|---|---|
| Number of clans | 8 |
| Age of cubs that count towards reproduction | 6 months |
| Cub sex ratio at 6 months old | 0.5 |
| Male age of sexual maturity | 24 months |
| Female age of sexual maturity | 24 months |
| Maximum lifespan | 25 years |

##### S1.1.3 Process overview and scheduling

The model proceeds in monthly time steps. There are three major demographic phases processed in each month in the following order: monthly survival, male dispersal, reproduction. Each of these demographic process is applied, in turn, to all clans. The order in which clans are considered for the dispersal process is randomised at each simulation step. A flow-chart of model phases is shown in Fig. ED1.

##### S1.1.4 Design concepts

**Emergence** Population dynamics and carrying capacity emerge from the behaviour of the individuals. The individual life cycle and behaviour are implemented using either stochastic predictions from statistical models fitted to real world data or empirical patterns derived from field observation of spotted hyenas. Fitness-seeking behaviour is not explicitly modelled but is included implicitly in the life cycle and behaviours.

**Sensing** Dispersing males select a new clan with a probability proportionate to the relative number of young females present ($>$ 12 months & $\leq$ 60 months; Höner et al., 2007). Dispersing males are therefore assumed to know the number of young females in all eight clans at the time of dispersal. Reproductive females choose males based on observed female mate choice patterns, where females prefer males that were born into or immigrated into the clan after the female's birth. Females are therefore assumed to know their own birth date and the date of birth and/or arrival of all males in the clan. All individuals are assumed to know their age, sex, and social rank to apply appropriate survival, reproduction, and dispersal probabilities.

**Interaction** Multiple interactions exist in the model. Survival, reproduction, and twinning are all assumed to experience density dependence in response to clan size. Reproduction of higher ranked females affects the rank of all lower ranked individuals because new cubs gain a rank just below that of their mother, thus pushing others down the hierarchy. Reproduction and female survival affect male dispersal behaviour by changing the number of young females in a clan. Survival of individuals affects the rank of all lower ranked individuals as the rank of a deceased individual is filled by the individual below it.

**Stochasticity** All demographic and behavioural outcomes are considered to be probabilistic, which incorporates expected demographic stochasticity.

**Observation** The simulation returns clan and population-level summaries in each time step. These include information on clan and population size, sex ratio, age ratio, rates of survival, reproduction, and dispersal.

##### S1.1.5 Initialization

At initialization, all clans are assumed to contain the same individuals that were observed in Ngorongoro Crater at the beginning of the study (1996-04-12). The starting population composition is available as supplementary data (see section Data availability in main text).

##### S1.1.6 Input

SHIM does not take any explicit environmental inputs; however, it does take an input 'year', which is used as a random factor in all statistical models to account for inter-annual differences in both the biotic and abiotic environment. Providing 'year' as an input in SHIM runs the simulation under the assumption that biotic and abiotic environmental conditions observed in the given year remain constant over time.

##### S1.1.7 Submodels

All statistical models in SHIM (survival, reproduction, twinning, dispersal) were fitted using observational data on spotted hyenas in Ngorongoro Crater. All vital rates were modelled as binomial outcomes and so the model structure can be described in the same way: $P(\text{event}) =$

$g^{-1}(\mathbf{X}\beta + \mathbf{Z}b)$, where $g^{-1}$ is the logit link function, $\mathbf{X}$ is a matrix of fixed-effect regressors, $\beta$ represents a column-vector of the model coefficients, $\mathbf{Z}$ is a matrix of dummy variables representing each year-clan combination and $b$ is a column-vector of the estimated random effect values. Random effects are not included in the model used for male additional dispersal, so in this particular model $\mathbf{Z}b = 0$.

All vital rate models are generalised linear mixed effects models with a binomial response, fitted using the R package spaMM (Rousset and Ferdy, 2014). Models include polynomial effects of age to allow for complex age relationships. The polynomial orders used for each model were chosen using both comparison of goodness of fit between nested models based on likelihood ratio tests and visual inspection of modelled age relationships. Visual inspection excluded any polynomials for which the direction of the age effect changed more than once over an individuals lifetime (i.e., first derivative of age cannot cross zero more than once). Such complex relationships were considered biologically unrealistic.

Below we describe each statistical model in more details, including specific information about the predictor variables used.

**Common predictors** A number of terms occur frequently across multiple sub-models. Each of these common terms is described here. Further detail about the biological relevance of each term in corresponding models is included below in the model descriptions.

- 'age': Age of individual in months. The variable 'age' is also assumed to interact with 'rank' as we expect the rate of senescence to be more rapid for low ranking individuals.
- 'rank': A categorical variable indicating whether natal individuals are in the top, middle, or bottom third based on their position in the hierarchy. Note that for immigrant individuals (which are always males in the simulations), no information on ranks is required (see Survival).
- 'clan': A categorical variable indicating the current clan where an individual is found. Accounts for differences between clans that may result from differences in variables like territory size and quality. The current clan of an individual is constant for all females, but can change over time for males due to dispersal.
- 'clan size': The total number of individuals (including all age groups and sexes) in a clan. This term is used to model density dependent effects. The variable 'clan size' is also assumed to interact with 'rank' as we expect density dependent effects to be more detrimental to lower ranking individuals.
- 'year': The calendar year in which observations were made. This incorporates variation in environmental conditions over time that may affect vital rates (e.g., availability of food). All models but one (additional dispersal) do include a random effect of year nested within

clan, which allows for different patterns of inter-annual variation in vital rates between clans (e.g., due to localised outbreaks of disease).

**Reproduction** Reproduction exclusively impacts female spotted hyenas (i.e., male mate choice is not considered). All adult females ($\geq$ 24 months) have the possibility to reproduce. Reproduction is defined as a female giving birth to a litter with at least one cub surviving to six months old. The minimum inter-birth interval for females is fixed to be four months to account for the approximate 110 day gestation period in the species. Females select the male with the longest tenure (i.e., has been in the clan for the longest) from a set of suitable males that match observed female mate choice patterns (Höner et al., 2007). A male must be an adult ($\geq$ 24 months) and have been born in or immigrated into the clan after the female was born. When no suitable males exist, a female cannot reproduce in that month.

We fitted separate models for primiparous and non-primiparous female reproduction.

Both primiparous and non-primiparous models include the following predictors:

- 'rank': As described above in Common predictors.
- 'time since last reproduction (or time since birth)': This term, computed as the logged number of months (+ 0.01 to avoid -Inf), shapes the age at first reproduction and inter-birth interval of reproducing females. For primiparous females the predictor describes months since birth. For non-primiparous females, this term represents the duration since last reproduction. Although rare, pregnancy before weaning the previous litter has been observed in Ngorongoro Crater (see Fig. S3) and other populations of spotted hyena (Holekamp et al., 1996). We did not therefore place a hard limit on inter-birth interval and allowed for the possibility that females could reproduce again four months after giving birth (i.e., following 110 day gestation).
- 'clan': As described above in Common predictors.
- 'clan size': As described above in Common predictors.
- 'effort': Mean monthly observation effort for a given clan in the year after a reproduction outcome was assessed. We calculated this observation effort as the proportion of individuals of all ages known to be in the clan that were actually observed at least once during the month as a proxy for observation effort. Increased observation effort should increase the chance of observing young cubs, and should thus lead to more accurate assessment of reproduction outcomes and thus higher probability of reproduction. When vital rate models are used to predict outcomes in SHIM, the observation effort is fixed to 1.0 to mimic the absence of any bias caused by imperfect observations. Observation effort in a given month is calculated as the proportion of known individuals observed during the month.

- ‘time since last reproduction (or time since birth):rank’: Interaction between ‘time since last reproduction or birth’ and ‘rank’. We expect high ranking females to have earlier age at first reproduction and shorter inter-birth interval than lower ranking females.
- ‘clan size:rank’: Interaction between rank and clan size.
- ‘year:clan’: A categorical variable used to define the levels for the random effects. As described above in Common predictors.

In addition to the above variables, the non-primiparous model also includes the term:

- ‘age’: As described above in Common predictors.
- ‘last reproduction:age’: This term allows for inter-birth interval of females to change with age. We expect inter-birth interval to become longer as females age.

**Twinning** Females that reproduce, based on the outcome of the reproduction models described above in Reproduction, can produce either single-cub or multi-cub litters. We grouped observed cases with litter size of both two and three together, which allowed us to work with a binomial response variable. A female with a multi-cub litter has at least two cubs that survive to at least six months old. Because cases of three cub litters are rare in the wild, we refer to the probability of multi-cub litters as ’twinning’ throughout the manuscript for convenience.

The twinning model includes the following predictors:

- ‘age + $age^2$’: Linear and quadratic effect of age. This accounts for the fact that multi-cub litters are less common in younger and older individuals.
- ‘rank’: As described above in Common predictors.
- ‘clan’: As described above in Common predictors.
- ‘clan size’: As described above in Common predictors.
- ‘effort’: As described in Reproduction. Mean monthly observation effort for a given clan in the year after a litter size was assessed. Increased observation effort should increase the chance of observing young cubs, and should thus lead to more accurate assessment of litter size and higher twinning probabilities. Observation effort in a given month is calculated as the proportion of known individuals observed during the month.
- ‘(age + $age^2$):rank’: Interaction between (linear and quadratic) age and rank. Effects of age are expected to be less pronounced in higher ranking individuals.
- ‘clan size:rank’: As described above in Common predictors.

- 'year:clan': A categorical variable used to define the levels for the random effects. As described above in Common predictors.

Due to the rarity of three cub litters all multi-cub litters are given a litter size of two within SHIM.

**Survival** Due to life-history differences between male and female spotted hyenas, we fitted separate male and female survival models. These are each described below.

Survival of all females greater than or equal to six months was modelled in a single model. All individuals younger than six months were excluded as early life survival is already modelled in the reproduction and twinning models described above in Reproduction and Twinning.

The female survival model includes the following predictors:

- 'age + age$^2$': Quadratic effect of age. This accounts for the fact that survival is lower in younger and older individuals.
- 'rank': As described above in Common predictors.
- 'clan': As described above in Common predictors.
- 'clan size': As described above in Common predictors.
- '(age + age$^2$):rank': Interaction between (quadratic) age and 'rank'. Effects of age are expected to be less pronounced in higher ranking individuals.
- 'clan size:rank': As described above in Common predictors.
- 'year:clan': A categorical variable used to define the levels for the random effects. As described above in Common predictors.

Male survival is modelled separately for juvenile males ($<$ 24 months) and adult males ($\geq$ 24 months). The structure of the juvenile male survival model is the same as that for female survival. In contrast, the survival model for adult males includes the following predictors:

- 'age + age$^2$': Quadratic effect of age. This accounts for the fact that survival is lower in younger and older individuals.
- 'post-dispersal status': Whether a male is in their birth clan (philopatric) or have dispersed to a new clan (disperser). Philopatric males maintain the rank in their birth clan, so we expect survival to be higher than dispersing males that are at the bottom of the new hierarchy. Effects of age are expected to be less pronounced in philopatric males.

- 'clan': As described above in Common predictors.
- 'clan size': As described above in Common predictors.
- '(age + age$^2$):post-dispersal status': Interaction between (quadratic) age and dispersal status.
- 'year:clan': A categorical variable used to define the levels for the random effects. described above in Common predictors.

Note that because the majority of adult males disperse (i.e., are not philopatric), most individuals are low ranking. Because of this, we did not include the term 'rank' in the survival model for adult males. The effect of post-dispersal status (philopatric or disperser) likely captures some of the effects of rank because philopatric males keep their position in the hierarchy of their birth clan.

**Dispersal** At 24 months, all males are considered to be adults and attempt to disperse. This primary dispersal is not determined by a statistical model and occurs for all male individuals once they become adults. The probability for a male to disperse to a given clan is considered proportional to the percentage of total young females in the population that are found in the clan. This also allows for the possibility of philopatric males that remain in their birth clan. Once a male has selected a clan for the first time (either philopatric or disperser), there is a possibility they will disperse again determined by a statistical model. Our fitted model of the probability of additional dispersal contains the following single predictor:

- 'age': As described above in Common predictors.
- 'post-dispersal status': Whether a male has never left their birth clan (philopatric) or not (disperser). We expect dispersal to be more common in philopatric males as they are expected to have fewer reproductive opportunities due to expected female mate choice behaviour (Höner et al., 2007).

#### S1.2 Pattern-oriented modelling

We used a pattern-oriented modelling approach (Grimm and Railsback, 2012) to examine the validity of our 'Spotted Hyena Individual-based Model' (SHIM). To test the patterns emerging from the simulation we ran an iteration of SHIM with the same statistical models and initial conditions as used for estimating $K_t$; however, rather than running the simulation until reaching a demographic equilibrium, the simulation was run for only 320 time-steps (months) to end in December 2022. The environmental conditions within SHIM were allowed to vary between years to mirror those observed in the real population. The realised value of the random effect in all statistical models was fixed to the corresponding calendar year in the simulation. In this way, the simulated data should cover similar length and inter-annual variation as the real data. Below we compare a number of different summary statistics between the simulation and real data.

##### S1.2.1 Lifespan

To compare the lifespan of individuals between simulated and real data we considered all uncensored individuals (i.e., individuals that were born and died during the observation period), which survived to at least six months old. Individuals that died before six months were excluded because this early life survival is incorporated into reproduction models. Mean lifespan of females in the simulation was 4.56 years (standard error = 0.13) while in the real data it was 4.78 years (standard error = 0.15). Mean lifespan of males in the simulation was 4.86 years (standard error = 0.13) compared to 4.36 years in the real data (standard error = 0.13). Maximum lifespan was slightly longer in the simulation (females: 20.08 years; males: 19.90 years) than in the real data (females: 18.45 years; males: 18.24 years). See also Fig. S1.

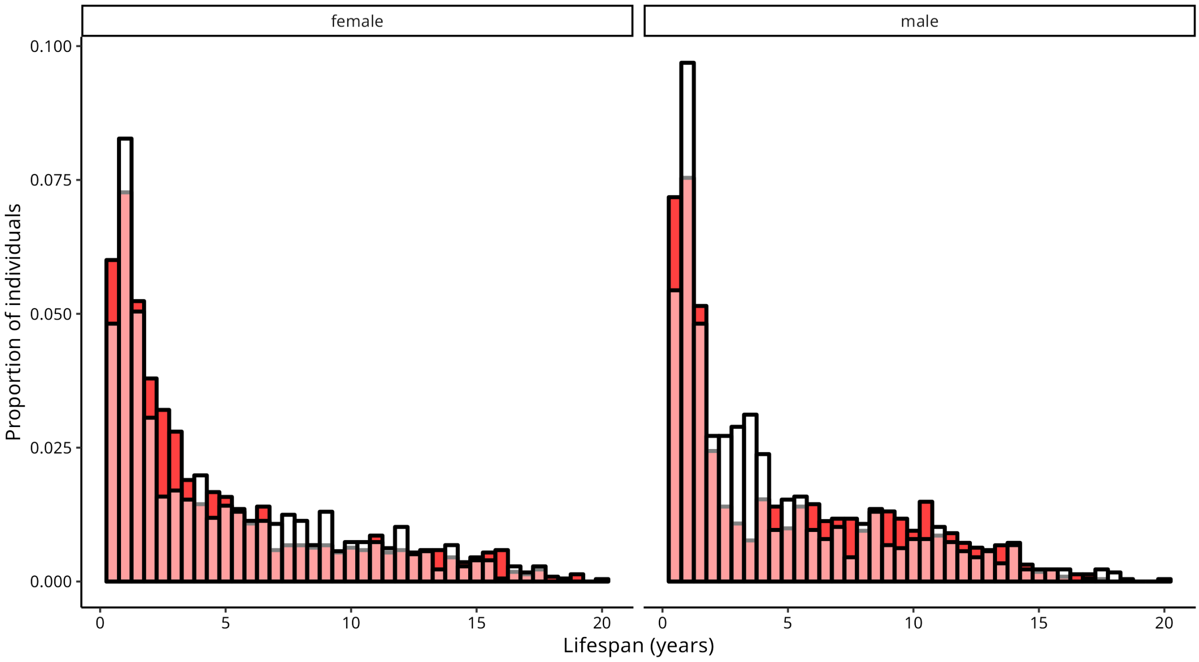


Figure S1: **Histogram showing the distribution of spotted hyena lifespans from SHIM (red) and observed data in Ngorongoro Crater (white).** Individuals that died before six months were excluded, as early life survival is incorporated into reproduction models. Plots only consider uncensored individuals that were born and died during the observation period.

##### S1.2.2 Age at first reproduction

To assess the observed and simulated distribution of the age at first reproduction we identified all females that were not left censored (i.e., born during the observation period) and had given birth to at least one litter where at least one cub had survived $\geq$ six months. Mean age at first reproduction of females was almost identical between the simulation (4.26 years; standard error = 0.08) and the real data (4.25 years; standard error = 0.07). See also Fig. S2.

##### S1.2.3 Inter-birth interval

To assess inter-birth interval, we again considered only reproductive events that produced at least one cub that survived $\geq$ six months. In this case, we included left censored adult females but only considered the interval between reproductive events that occurred during the study period. Inter-birth interval of females was slightly shorter in the simulation (16 months; standard error = 0.22) than the real data (20 months; standard error = 0.27). There was an over-representation of low values in the simulated data. See also Fig. S3.

##### S1.2.4 Lifetime reproductive success

Lifetime reproductive success of all uncensored adult females was calculated for both the simulated and real data. As with other measures of reproduction, we only considered cubs that survived $\geq$ six months. Although females reproduced slightly too early in the simulation, their overall lifetime reproductive success was very similar between SHIM and real data. Mean lifetime reproductive success of females in the simulated data was 2.81 (standard error = 0.14) while in the real data it was 2.70 (standard error = 0.13). Maximum lifetime reproductive success was also similar between the simulation (18 offspring) and real data (17 offspring). See also Fig. S4.

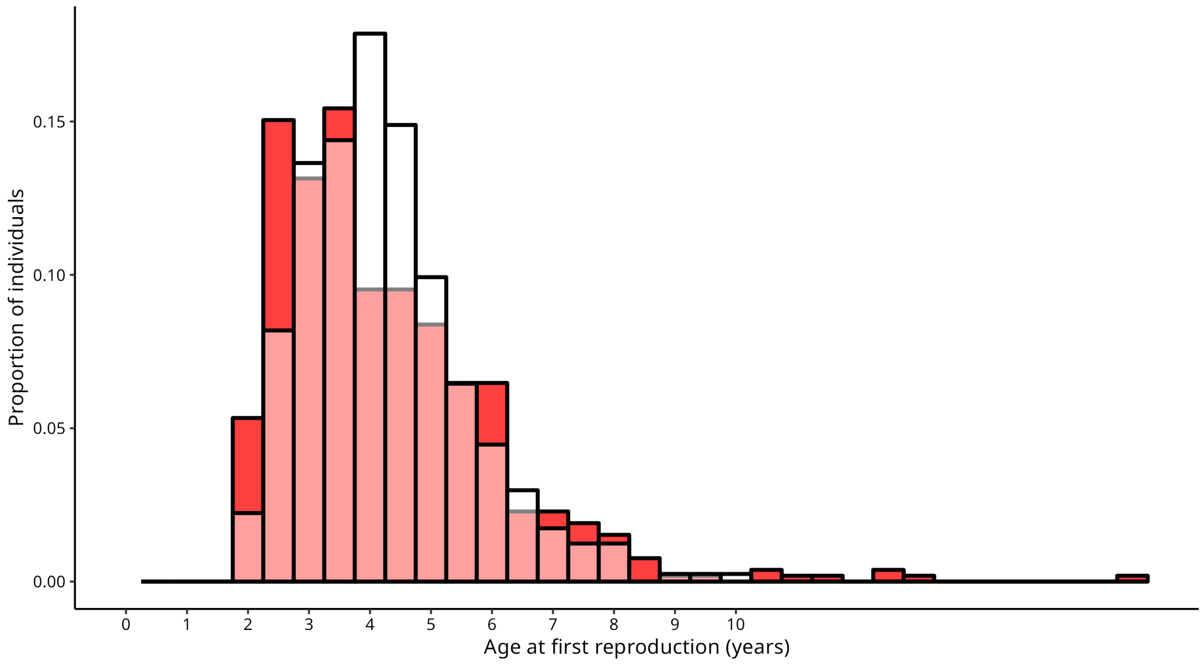


Figure S2: **Histogram showing the distribution of age at first reproduction for female spotted hyenas from SHIM (red) and observed data in Ngorongoro Crater (white)**. Data only considers females that were not left censored and reproductive events where at least one cub had survived six months or more.

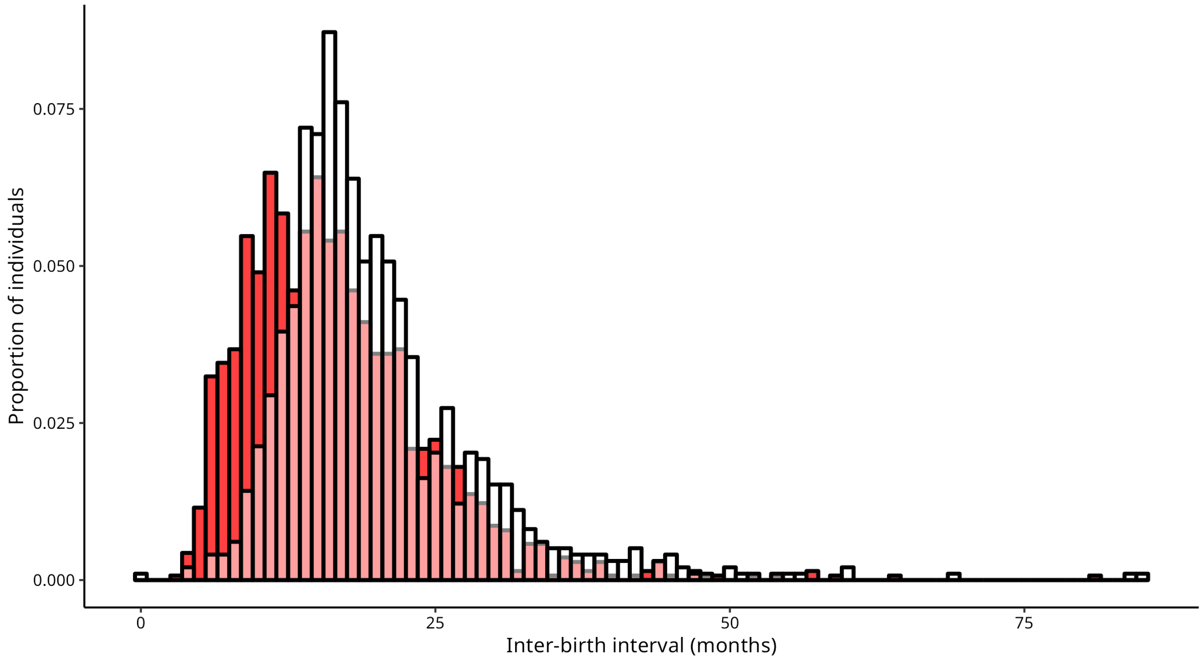


Figure S3: **Histogram showing the distribution of inter-birth intervals for female spotted hyenas from SHIM (red) and observed data in Ngorongoro Crater (white).** Only considers reproductive events where at least one cub had survived six months or more.

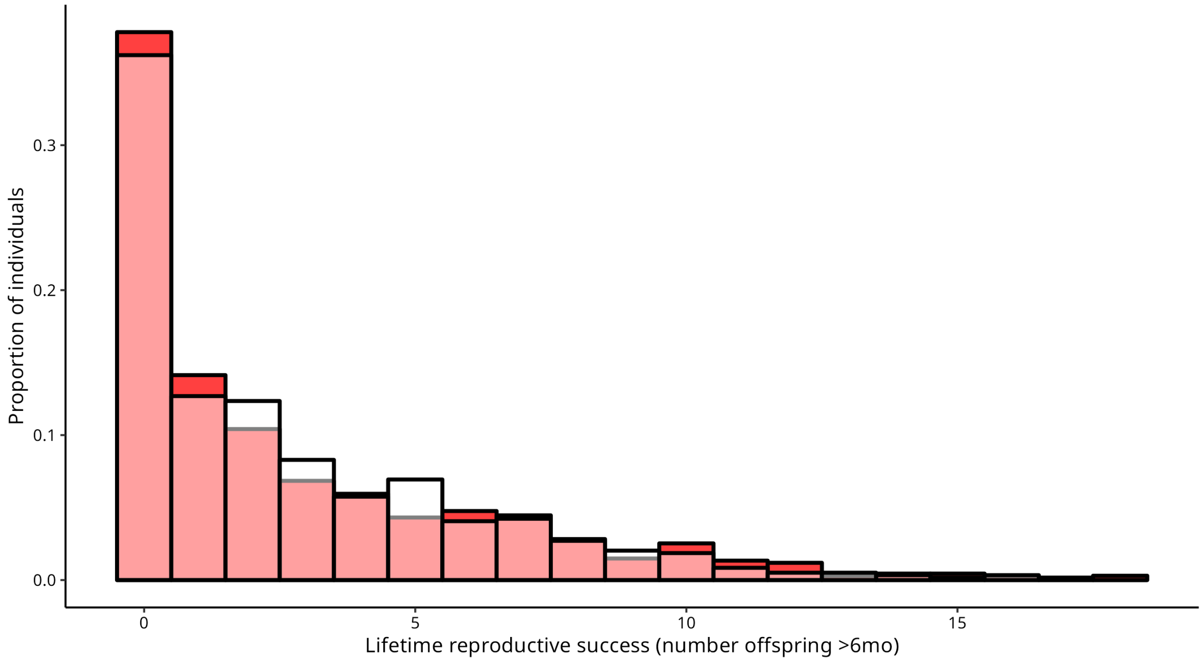


Figure S4: **Histogram showing lifetime reproductive success of female spotted hyenas from SHIM (red) and observed data in Ngorongoro Crater (white).** Only considers uncensored females and cubs that survived six months or more.

##### S1.2.5 Effects of rank

To compare effects of rank between simulated and real data we compared lifetime reproductive success (described above) between females from the three rank categories. In both simulated and real data, lifetime reproductive success was higher in the higher rank categories. The simulation tended to slightly underestimate lifetime reproductive success of the top ranking individuals and slightly overestimate lifetime reproductive success of middle ranking individuals (Fig. S5). Lifetime reproductive success of bottom ranked females was similar between the simulation and observed data.

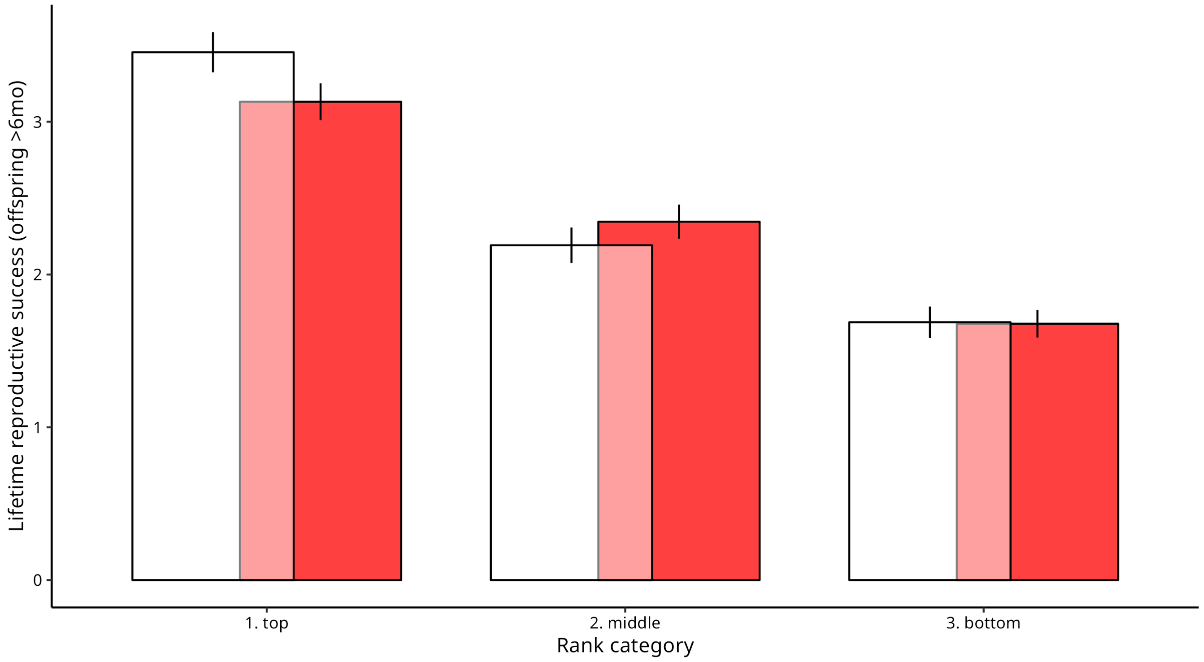


Figure S5: **Histogram showing lifetime reproductive success of female spotted hyenas at different ranks from both SHIM (red) and observed data in Ngorongoro Crater (white)**. We assigned female rank categories based on the rank of females among all natal individuals (i.e., excluding immigrant males) at the point when the female first reached sexual maturity (i.e., 24 months).

### S2 Supplementary Material 2: Additional results

#### S2.1 Trends in age- and sex-ratios

The age ratio of the population was strongly biased against juveniles ($\leq$ 24 months) for the full study period (median annual ratio juveniles/all individuals = 0.434; range = 0.375–0.476; red line in Fig. 2b). Bias against juveniles was stable during the study period ($\beta_{\text{ageratio}}$ = -0.00278; $\text{CI}_{95\%}$ = -0.00976/0.00544; Likelihood Ratio Test: $\chi^2$ = 0.646; df = 1; $p$ = 0.42). Adult sex ratio (number of adult males/total number of adults in a given year) showed a slight female bias for the majority of the study period, ranging from 0.436 to 0.521 (median = 0.472, black line; Fig. 2b). Adult sex ratio showed no significant change over time ($\beta_{\text{sexratio}}$ = -0.00323; $\text{CI}_{95\%}$ = -0.00964/0.00388; Likelihood Ratio Test: $\chi^2$ = 1.03; df = 1; $p$ = 0.31).

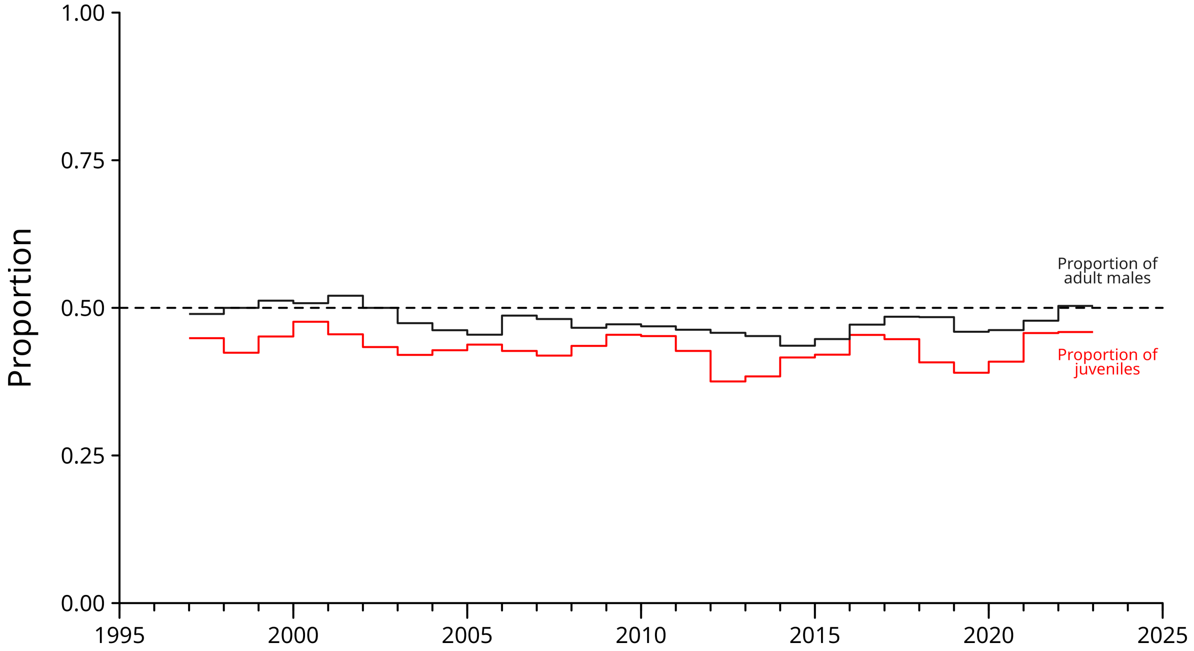


Figure S6: **Proportion of adult males (adult males/all adults; black line), and proportion of juvenile (juveniles/all individuals; red line) spotted hyenas in Ngorongoro Crater, Tanzania since 1996**. Dashed line represents a value of 0.5 at which there would be no bias towards a particular age group or sex.

#### S2.2 Vital rates, density dependence, and carrying capacity

In our individual-based model, $K_t$ is an emergent property of the expression of the vital rates of all individuals in the population. We here detail the link between the vital rates and carrying capacity, and their relation to density dependence. Simulation runs stabilized around a fixed population size ($K_t$) because of the existence of density dependence in vital rates. We included density dependence in six out of the seven statistical models that shape simulation outcomes (Table ED2; section S1.1.7). Specifically, we included density dependence in the models predicting female survival, juvenile male survival, adult male survival, female primiparous reproduction, female non-primiparous reproduction, and twinning (Fig. ED2). Density dependence was not included in the model used to predict dispersal of males beyond their first clan selection, which is a rare event.

We considered the strength of density dependence (i.e., slope of clan size effect) to be affected by the rank of the individuals (Fig. ED2; Fig. S7; Fig. S8; Table ED2) since social rank strongly impacts competition for resources in spotted hyenas (Frank, 1986; Höner et al., 2010). Models accounted for other confounding variables such as age, but the effects of these confounding variables was considered to be independent of density dependence. We therefore did not consider year to affect the strength of density dependence directly. However, since the nested random intercept term (year within clan) influenced the simulated life history outcomes—which translated into different clan size—predictions of $K_{\mathrm{t}}$ differed from year to year, and from clan to clan, due to changing environmental conditions affecting all six vital rates.

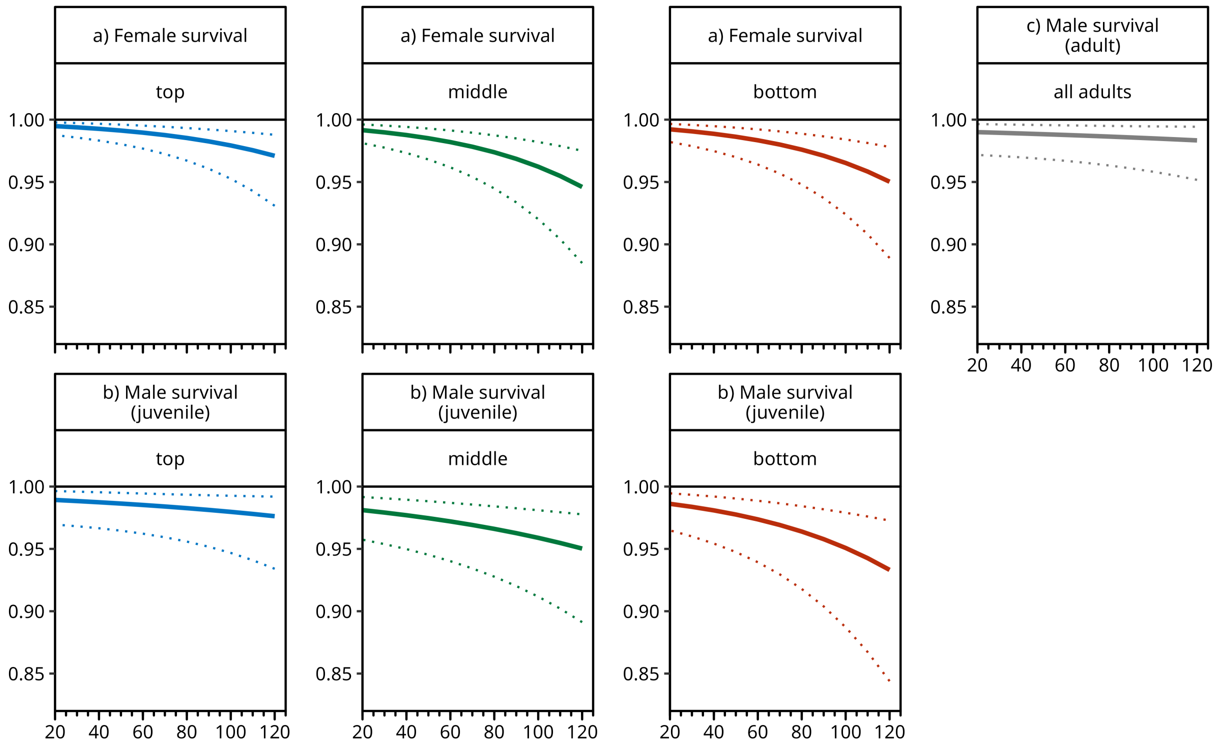


Figure S7: **Density dependence in survival models.** This figure is a replicate of Fig. ED2 which depicts all social rank categories separately for all survival models.

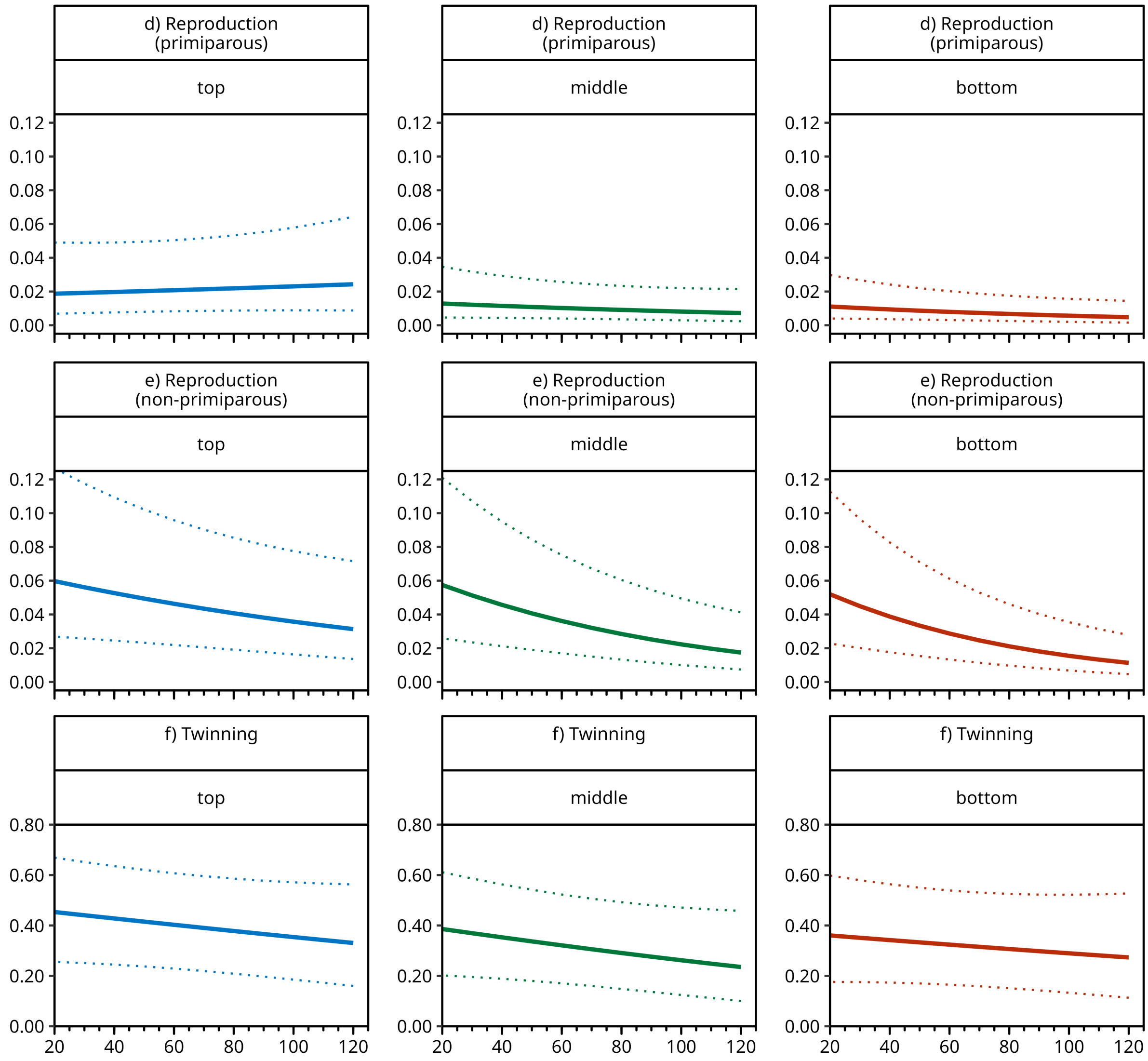


Figure S8: **Density dependence in reproduction and twinning models.** This figure is a replicate of Fig. ED2 which depicts all social rank categories separately for all reproduction and twinning models.

Given that SHIM incorporates density dependence only at the level of clan, it is worth inspecting density dependence at the population level as well. For this, we used the ten simulation runs for an example year (2001) and visualised the relationship between population size at time t ($N_t$) and population growth ($\lambda$) as $\frac{N_{t+1}}{N_t}$. Fig. S9 shows a clear negative relationship between population size ($N_t$) and population growth ($\lambda$) at the level of the population.

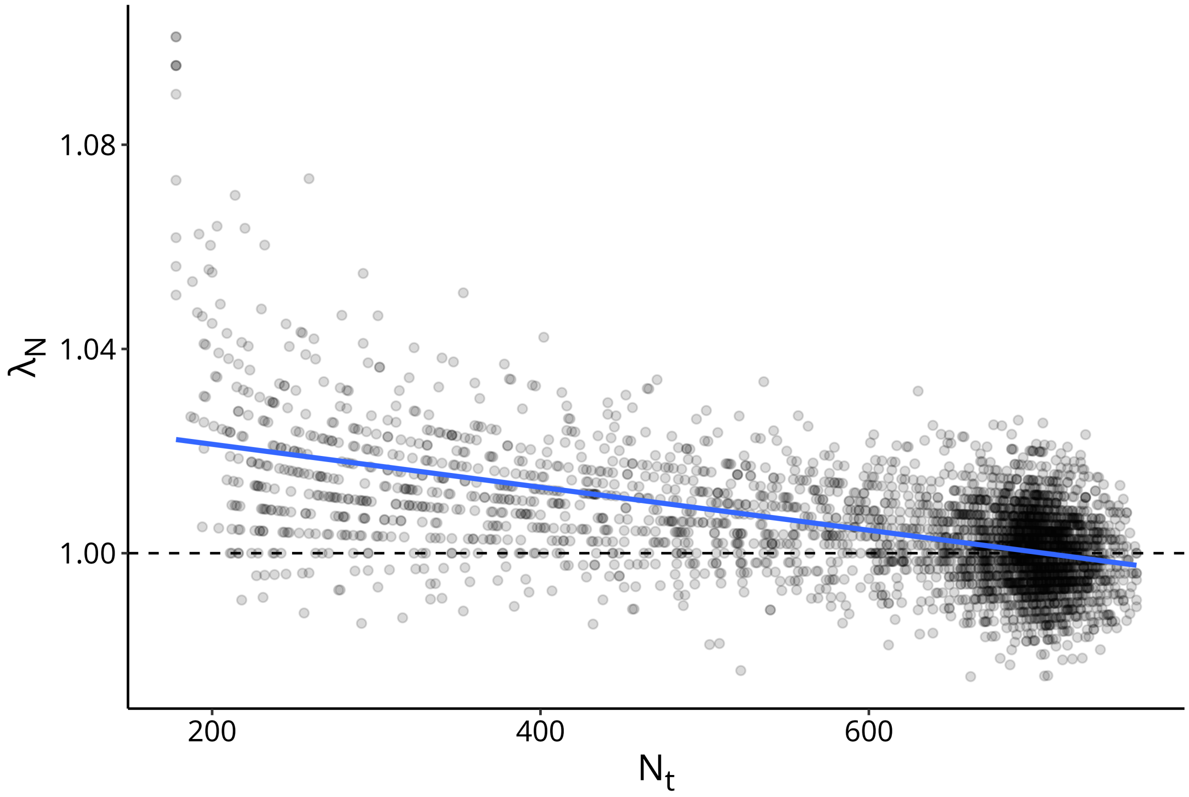


Figure S9: **Density dependence at the level of the population.** Population growth $\lambda_{\rm N}$ decreases with increasing population size $N_{\rm t}$, demonstrating a negative density dependent relationship. Blue line shows a simple linear model fitted through the data.

#### S2.3 Estimation of carrying capacity when assumed static

This section provides additional details of results presented in Results section 2.2. When simulations were no longer considered to be conditional on the year random effect estimate, SHIM produced a fixed carrying capacity of 547 individuals (range: 532–553), which is 94-98% of the median value for $K_{\rm t}$ calculated between 1997 and 2022. Estimated fixed carrying capacity using Ricker and Beverton-Holt equations were similar, although slightly lower (range: 526–533 (Fig. S10). The latter estimates were obtained considering both the exponential and hyperbolic versions of the Ricker and Beverton-Holt equations (Johst et al., 2008). To compute them, we used the observed population growth in Ngorongoro measured over six-month time steps between 1996 (July–December) and 2022 (July–December) as inputs. We jointly estimated carrying capacity and maximum growth rate ($R_{\rm max}$) in each equation using the BOBYQA optimization algorithm from the nloptr package in R, which we found more successful at minimizing the mean squared error between observed and predicted population growth between time $t$ and $t+1$ than the alternatives we tried (COBYLA, LBFGS, PRAXIS, Nelder-Mead, and SBPLX). To account for sensitivity of the optimization to initial parameter values we ran each algorithm using a range of initial parameter values and used the estimate of carrying capacity that returned the lowest mean squared error value.

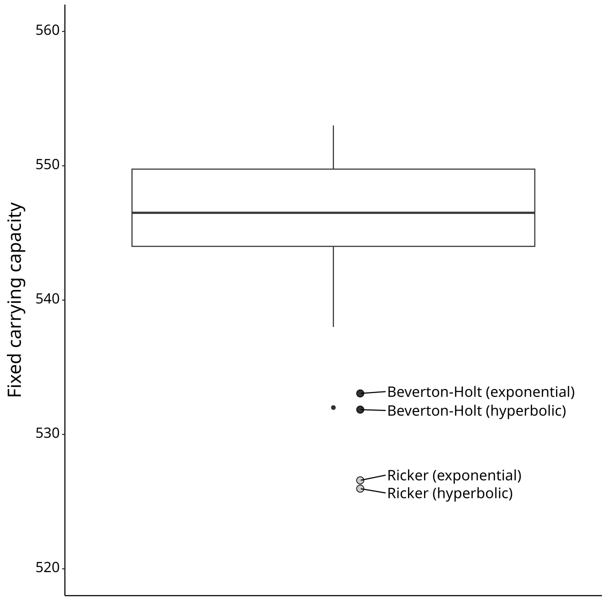


Figure S10: **Estimated fixed carrying capacity is similar using SHIM and traditional demographic models.** Filled points show estimate of carrying capacity according to exponential and hyperbolic Ricker and Beverton-Holt equations. The boxplot shows fixed carrying capacity as estimated using SHIM (547 individuals; range: 532–553). The boxplot is traditionally representing the median (thick horizontal line), inter-quartile range (box), with whiskers reaching all points that are no further away from the hinge of the box than 1.5 times the interquartile range. There is one points beyond this limit shown as a small point located left of the Beverton-Holt hyperbolic estimate.

#### S2.4 Change in carrying capacity over time

This section provides additional details of results presented in Results section 2.2. There was no significant change in carrying capacity over time in either the full study period ($\beta_{\text{year}}$ = -5.24 individuals per year; $\text{CI}_{95\%}$: -11.9/1.62; Likelihood Ratio Test: $\chi^2$ = 2.21; df = 1; $p$ = 0.14) or the period of population growth up to 2011 ($\beta_{\text{year}}$ = 14.6 individuals per year; $\text{CI}_{95\%}$: -2.03/30.3; Likelihood Ratio Test: $\chi^2$ = 3.23; df = 1; $p$ = 0.072). At the level of clan, six of eight clans showed no significant change in clan level carrying capacity ($K_{\text{tc}}$) over the study period (Table S2). In one clan (Lemala), there was a significant negative change in clan level carrying capacity over time (Table S2). Only one of the eight clans (Ngoitokitok) showed significant changes in population size when looking solely at the period of population growth (Table S3). AR1 temporal auto-correlation in $K_{\text{tc}}$ ($\phi$) was 0.62 for the full study period and 0.39 for the period of population growth (1997–2011).

Table S2: **Slope and 95% confidence interval of carrying capacity change over time at the level of clan ($K_{tc}$) for all eight spotted hyena clans of Ngorongoro Crater during the full study period (1997–2022).**

| Clan | $\beta_{year}$ | $CI_{95\%}$ | |
|---|---|---|---|
| Airstrip | 0.507 | -1.43 | 2.55 |
| Engitati | -1.10 | -3.04 | 0.940 |
| Forest | -0.190 | -2.12 | 1.85 |
| Lemala | -2.57 | -4.51 | -0.526 |
| Munge | -1.15 | -3.09 | -0.893 |
| Ngoitokitok | -0.889 | -2.82 | 1.15 |
| Shamba | 0.300 | -1.63 | 2.34 |
| Triangle | 0.319 | -1.62 | 2.36 |

Table S3: **Slope and 95% confidence interval of carrying capacity change over time at the level of clan ($K_{tc}$) for all eight spotted hyena clans of Ngorongoro Crater during the period of initial population growth (1997–2011).**

| Clan | $\beta_{year}$ | $CI_{95\%}$ | |
|---|---|---|---|
| Airstrip | 2.29 | -0.409 | 4.84 |
| Engitati | 0.725 | -1.90 | 3.35 |
| Forest | 2.10 | -0.531 | 4.72 |
| Lemala | 0.882 | -1.74 | 3.50 |
| Munge | 2.32 | -0.306 | 4.94 |
| Ngoitokitok | 2.81 | 0.187 | 5.44 |
| Shamba | 1.40 | -1.23 | 4.02 |
| Triangle | 2.16 | -0.463 | 4.79 |

#### S2.5 Relationship between N and K

This section provides additional details of results presented in Results section 2.3. The annual population growth rate was strongly correlated with the distance between population size and carrying capacity at time t (i.e., how far the population was from (non-zero) demographic equilibrium). Fig. ED3 & S11b shows how the distance between carrying capacity (estimated from our model) and population size (directly observed) is related to population change in each year. All cases of annual population growth occurred when population size was below carrying capacity (left side; Fig. ED3 & S11b). More specifically, population growth was generally higher in years where population size was further below carrying capacity, with a strong correlation between $\lambda_N$ and $K_t - N_t$ (Pearson's r = 0.84), much higher than that observed between $\lambda_N$ and $\lambda_K$ (r = 0.22). In those few cases where population size *exceeded* carrying capacity, we observe clear population decline, as would be expected from theory (right side; Fig. ED3 & S11b). In 2019, for example, we can see that population size was above carrying capacity (Fig. ED3 & S11b, red point), which led to a decline in population size despite increasing carrying capacity during the year (Fig. 4 & S11a).

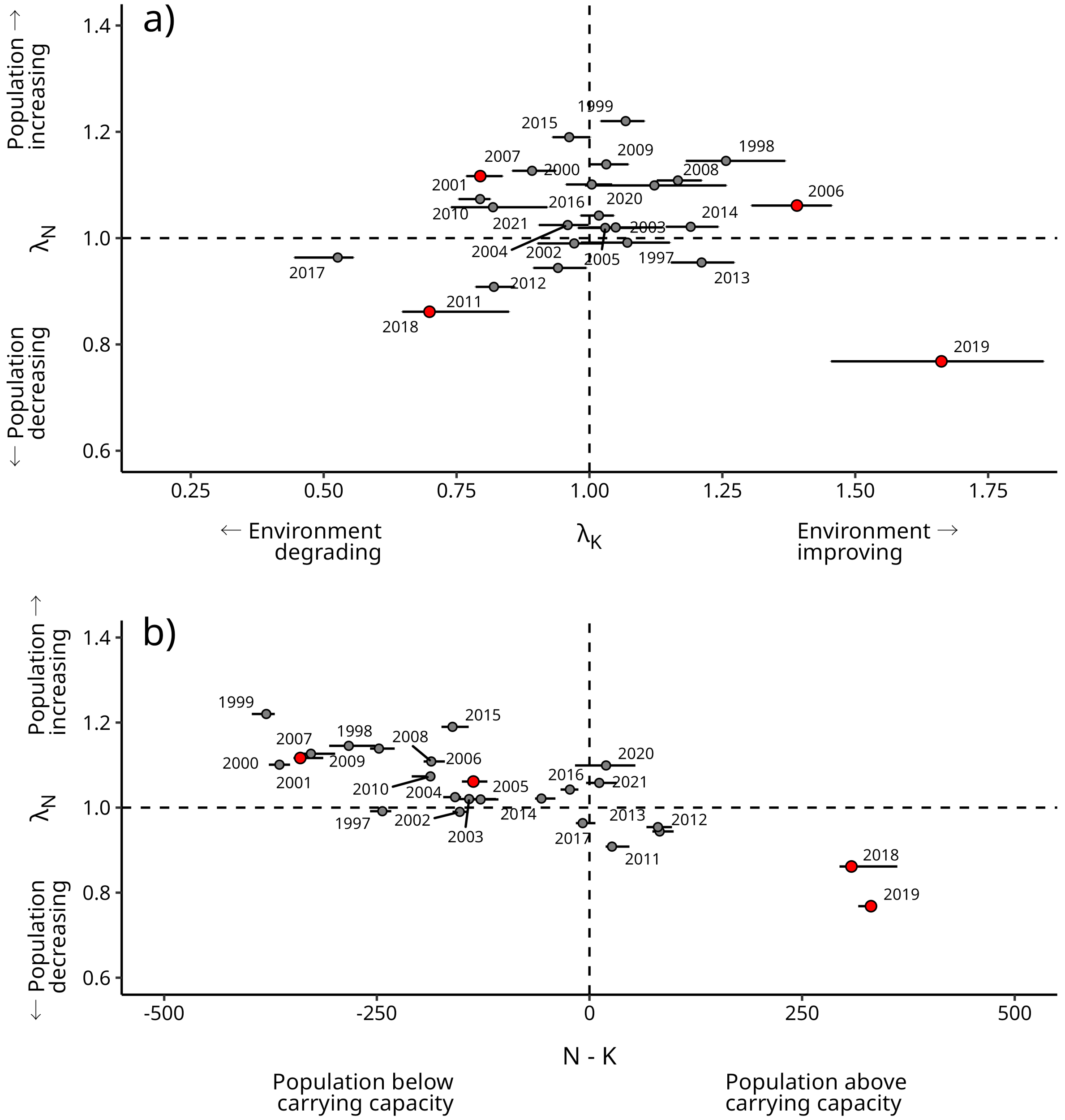


Figure S11: **Inter-annual change in population size ($\lambda_N$) is uncorrelated with inter-annual change in time-varying carrying capacity ($\lambda_K$) in spotted hyenas but is but strongly correlated with $N_t - K_t$.** a) Replication of Fig. 4 including year labels. b) Replication of Fig. ED3 including year labels. For visual clarity only the first year of each period is indicated in the plot. For example, 1997 corresponds to values for 1997–1998.

#### S2.6 Variation in vital rates over time

This section provides additional details of results presented in Results section 2.4. We found no significant change in survival over time, while there was a significant *decline* in both primiparous and non-primiparous reproduction and twinning (Table S4). Figure S12 includes partial dependence plots showing changes in vital rates over time. Table S4 shows outcome of likelihood ratio test comparison between models with and without year and estimated coefficient of year in each model. Year coefficient is presented on the scale of the linear predictor used in the models (logit link).

Table S4: **Vital rates have not increased over time**. Likelihood ratio test results comparing vital rates models with and without a quantitative year term. Model coefficients are expressed on the scale of the linear predictor.

| Response variable | $\beta_{\text{year}}$ | $\chi^2$ | df | $p$ |
|---|---|---|---|---|
| Female survival | -0.00678 | 0.675 | 1 | 0.41 |
| Male pre-dispersal survival | -0.00158 | 0.0432 | 1 | 0.83 |
| Male post-dispersal survival | -0.00917 | 0.585 | 1 | 0.44 |
| Twinning | -0.0736 | 30.8 | 1 | $< 0.001$ |
| Primiparous reproduction | -0.0885 | 51.4 | 1 | $< 0.001$ |
| Non-primiparous reproduction | -0.0823 | 66.9 | 1 | $< 0.001$ |

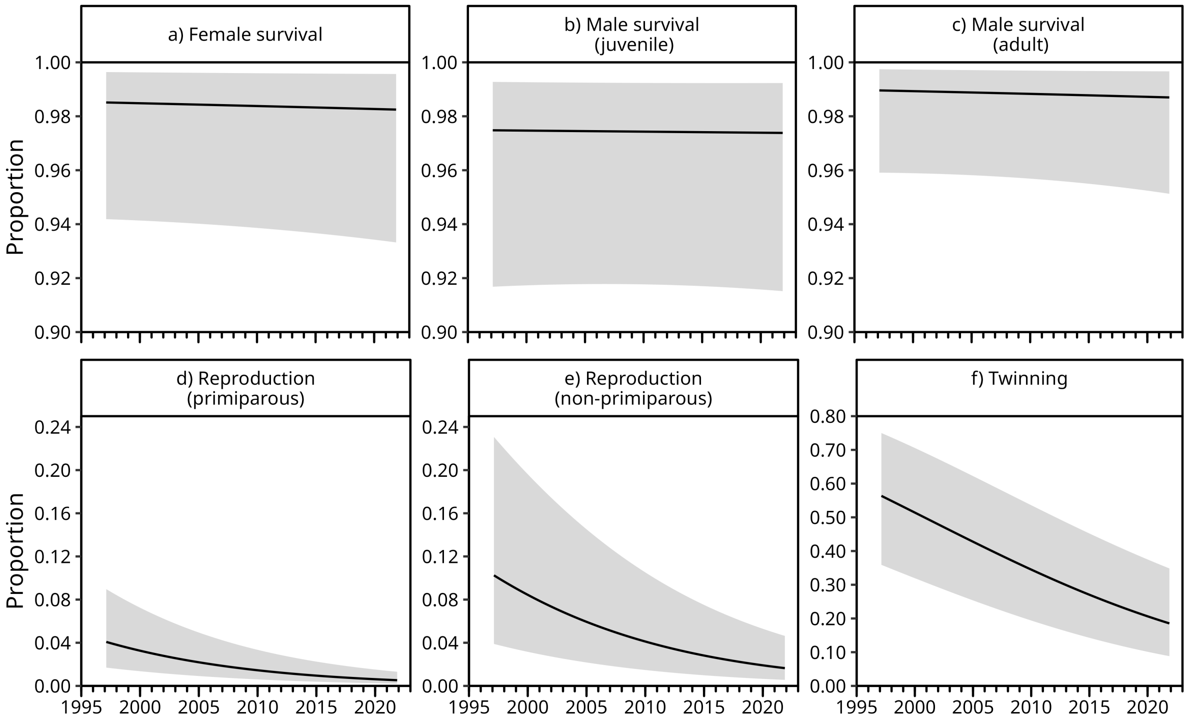


Figure S12: **Vital rates have not increased over time**. Plotted predictions show changes in vital rates given changes over time (year), computed as partial dependence effects (see caption Fig. ED2 for details). Note that y-axis limits differ between panels.

To understand the impact of environmental conditions on the vital rates of individuals we also tested for linear time-trends in each vital rate separately. We assessed the significance of an additional (quantitative) year predictor added a posteriori to each of the six vital rate models. Our results show that monthly probability of female survival, male survival did not present significant linear time-trend (Fig. ED4; Fig. S12). The probability of primiparous and non-primiparous reproduction and twinning did significantly *decrease* over time after accounting for confounding variables (Fig. ED4; Fig. S12), which is contrary to the trend observed for $N_t$. In agreement with results obtained for $K_t$ (Fig. 3), the recovery of the population of spotted hyenas since 1996 cannot thus be explained by changes in any particular vital rates.

Different vital rates impacted $K_t$ differently in our individual-based model. An elasticity analysis measuring the response of $K_t$ to proportional changes in each vital rate revealed that $K_t$ was much more sensitive to changes in female survival than to any of the other vital rates (section S2.7; Fig. S13).

To test for temporal trends in vital rates we extended models used inside SHIM (described in S1.1.7; Table ED2) by including a fixed effect of year modelled as a quantitative variable. These models allowed us to test for temporal trends while accounting for the effect of known covariates, such as age and clan, also impacting vital rates.

#### S2.7 Elasticity analysis

For each vital rate i, we calculated the elasticity of the time-varying carrying capacity ($E_{K_{ti}}$) as the change in absolute time-varying carrying capacity ($\Delta K_t$) given a consistent shift in vital rates. We modified individual statistical models used inside SHIM to change (increase or decrease) vital rates and then recalculated $K_t$ in each year using the same methods as described above. Vital rate models were adjusted by shifting the intercept term, leaving all other estimates, and thus the effect of all co-variates in the vital rate models (e.g., age, social rank), the same. The intercept of each model was shifted to achieve a 25% change in odds (i.e., odds ratio of 1.25) considering the mean vital rate value conditional on the observed characteristics of all observed individuals used to fit the model.

Time-varying carrying capacity ($K_t$) was most affected by changes in female survival, particularly in earlier years S13. A 25% change in the odds of female survival altered estimates of $K_t$ between 70 and 238, depending on the year. In comparison, similar changes in odds for other vital rates led to at most a 43 individual change in $K_t$.

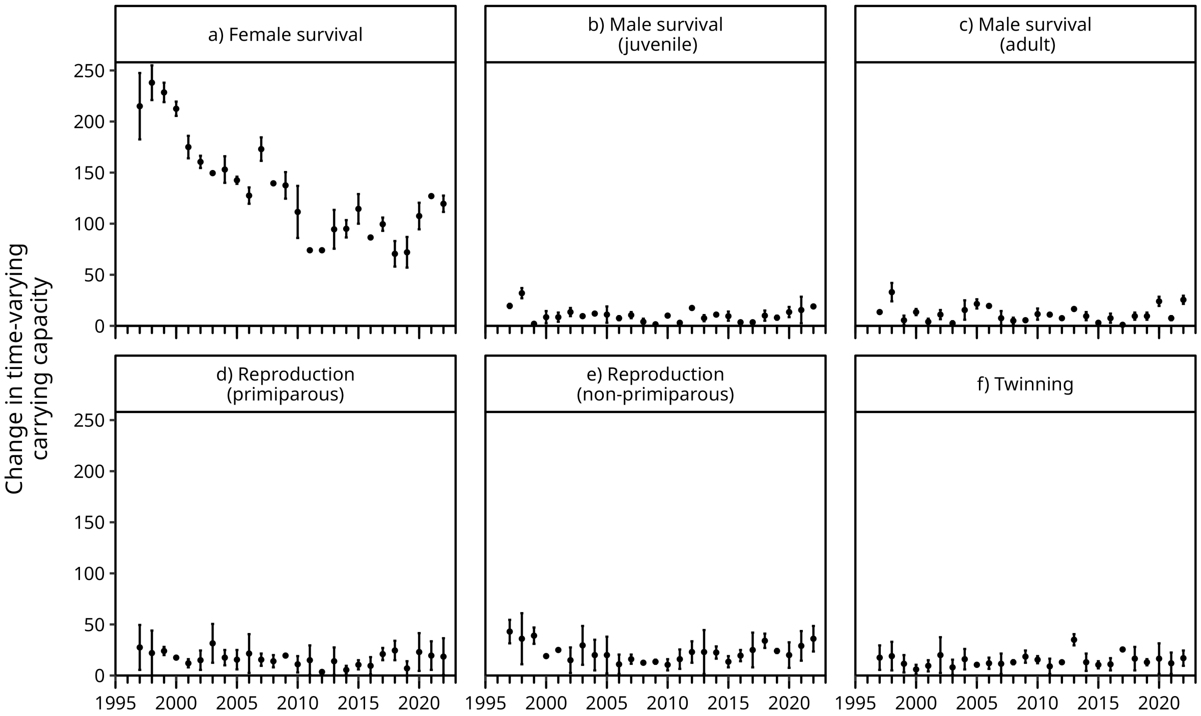


Figure S13: **Change in female survival has the strongest effect on time-varying carrying capacity.** Plot shows absolute change in time-varying carrying capacity in response to a change in vital rates equivalent to an odds ratio of 1.25.

#### S2.8 Biotic drivers of carrying capacity

This section provides additional details of results presented in Results section 2.4. Effects of per capita disease prevalence, prey abundance, and competitor density (lions) had a significant effect on clan carrying capacity in spotted hyenas. The effect of lion density and disease (both negative) and prey abundance (positive) matched our biological expectations (Table S5; Fig. 5). There was no evidence of effects of road use or grazing restrictions.

Table S5: **Table showing model coefficients for different potential drivers of carrying capacity in spotted hyena clans.** This table provides the estimate values for each parameters and their associated conditional standard errors. Note that quantitative predictors ('prey_abundance', 'per_capita_disease', and 'per_capita_lion_density') are expressed as Z-scores. For 'year' the variance of the random effect is provided as an estimate and no conditional standard error is given. Temporal auto-correlation term ($\phi$) was estimated at 0.874. The number of rows used to fit the model was 196.

| Parameter | Estimate | Cond. SE |
|---|---|---|
| intercept ($\text{clan}_{\text{A}}$, $\text{road\_use}_{\text{active}}$, $\text{grazing}_{\text{absent}}$) | 117 | 16.9 |
| prey_abundance | 6.71 | 1.90 |
| per_capita_disease | -4.47 | 1.80 |
| per_capita_lion_density | -6.30 | 2.45 |
| $\text{road\_use}_{\text{inactive}}$ | -17.1 | 13.9 |
| $\text{grazing}_{\text{present}}$ | -16.2 | 8.89 |
| $\text{clan}_{\text{E}}$ | -48.4 | 9.18 |
| $\text{clan}_{\text{F}}$ | -35.7 | 6.28 |
| $\text{clan}_{\text{L}}$ | -41.5 | 9.07 |
| $\text{clan}_{\text{M}}$ | -31.0 | 9.60 |
| $\text{clan}_{\text{N}}$ | -56.1 | 9.50 |
| $\text{clan}_{\text{S}}$ | -26.6 | 9.43 |
| $\text{clan}_{\text{T}}$ | -77.4 | 9.84 |
| year (random factor) | 511 | |
| residual variance | 496 | |

### S3 Supplementary Material 3: Additional discussion

#### S3.1 Possible impacts of climate change

Temporal shifts in climatic conditions have already become evident in East Africa over the past century, with increasingly frequent heavy rain events contrasted with more frequent and severe droughts (Nicholson, 2017). Heavy rainfall can elevate disease risk by increasing the abundance of disease vectors, such as biting insects (Gallana et al., 2013; Chowell et al., 2019). Conversely, drought events can increase disease transmission by increasing aggregation of potential hosts around limited water resources (Gallana et al., 2013; Franz et al., 2018) and acting as an additional stressor on wildlife, making them more vulnerable to infection (Acevedo-Whitehouse and Duffus, 2009).

Changes in the prey community are also expected as a consequence of climate change (Moehlman et al., 2020). Long-term trends towards hotter and drier conditions will likely favour smaller-bodied herbivores, such as gazelle, over those larger-bodied herbivores that depend on bulk-forage, such as buffalo (Moehlman et al., 2020). At the same time, increased rainfall variability is expected to lead to increased variation in the abundance of many key prey species for spotted hyenas (Moehlman et al., 2020). Climate change is also expected to affect the availability of suitable forage for herbivores by facilitating the spread of unpalatable invasive weeds during heavy rain events (Ngondya and Munishi, 2021; Nyarobi et al., 2022). Spotted hyenas are flexible in both their diet and hunting strategies (Hofer and East, 1993; Höner et al., 2002; Yirga et al., 2012) which should allow them to adjust to shifts in prey communities. However, any changes to hunting behaviour that require spotted hyenas to forage over larger distances will likely reduce vital rates—particularly for individuals lower in the social hierarchy (Hofer and East, 1993)—which should impact carrying capacity.

#### S3.2 Alternative methods for estimating time-varying carrying capacity

In order to predict $K_t$, the only structural requirement is that the method can predict population size and reach demographic equilibrium when environmental conditions are stable. Each possible approach presents pros and cons, and certain approaches could complement individual-based modelling in providing additional insights about the population dynamics which are relevant for conservation. In particular, 'Integral Projection Models' (IPM; Easterling et al., 2000) can been used to decompose the variation in population growth rates, and thus perhaps in $K_t$, into demographic components (stochasticity, population structure. . . ) (Koons et al., 2016; Knape et al., 2023).
